# A Powerful Method for Pleiotropic Analysis under Composite Null Hypothesis Identifies Novel Shared Loci Between Type 2 Diabetes and Prostate Cancer

**DOI:** 10.1101/2020.04.11.037630

**Authors:** Debashree Ray, Nilanjan Chatterjee

## Abstract

There is increasing evidence that pleiotropy, the association of multiple traits with the same genetic variants/loci, is a very common phenomenon. Cross-phenotype association tests are often used to jointly analyze multiple traits from a GWAS. The underlying methods, however, are often designed to test the global null hypothesis that there is no association of a genetic variant with any of the traits, the rejection of which does not implicate pleiotropy. In this article, we propose a new statistical approach, PLACO, for specifically detecting pleiotropic loci between two traits by considering an underlying composite null hypothesis that a variant is associated with none or only one of the traits. We propose testing the null hypothesis based on the product of the Z-statistics of the SNPs across two studies and derive a null distribution of the test statistic in the form of a mixture distribution that allows for fractions of SNPs to be associated with none or only one of the traits. We borrow approaches from the statistical literature on mediation analysis that allow asymptotic approximation of the null distribution avoiding estimation of nuisance parameters related to mixture proportions and variance components. Simulation studies demonstrate that the proposed method can maintain type I error and can achieve major power gain over alternative simpler methods that are typically used for testing pleiotropy. PLACO allows correlation in summary statistics between studies that may arise due to sharing of controls between disease traits. Application of PLACO to publicly available summary data from two large case-control GWAS of Type 2 Diabetes and of Prostate Cancer implicated a number of novel shared genetic regions near *ZBTB38* (3q23), *RGS17* (6q25.3), *HAUS6* (9p22.1), *UBAP2* (9p13.3), *RAPSN* (11p11.2), *AKAP6* (14q12), *KNL1* (15q15) and *ZNF236* (18q23).

## Introduction

Years of genetic research on various complex human traits have implicated numerous genetic variants as risk factors for two or more traits. Pleiotropy, the phenomenon where a genetic region or locus confers risk to more than one trait^1^, is widely observed for many diseases and traits^2^, especially cancers^3^, autoimmune^4^ and psychiatric^5,6^ disorders. It has also been observed in seemingly unrelated traits; for instance, early-onset androgenetic alopecia and Parkinson’s disease^7^, Crohn’s disease and Parkinson’s disease^8^, and coronary artery disease and tonsillectomy^9^. Pleiotropy provides new opportunities, as well as challenges, for diagnosis, therapeutics, and intervention on diseases^1,2,10,11^. Consequently, it is important to identify and study shared genetic basis of complex traits.

To detect potential pleiotropic effects of genetic variants, many statistical methods for jointly analyzing multiple traits have been proposed^1,12,13^. Use of these methods – commonly referred to as “cross-phenotype association tests” – has been gaining traction over the past few years, and has led to successful discovery and replication of genetic overlap among different human disorders and traits^5,14–21^. Typical cross-phenotype association methods test the global null hypothesis that no trait is associated with a given genetic variant against the alternative hypothesis that at least one of the traits is associated. Thus, rejection of the null hypothesis could just be due to one trait being associated with the genetic variant, and not necessarily due to pleiotropy.

Recently, a few methods have been proposed to specifically test for pleiotropy, where the rejection of the null hypothesis is driven by significant association of the genetic variant with more than one trait^22–25^. All of these methods require individual-level phenotype and genotype data on the same set of randomly sampled individuals, and cannot be readily extended to diseases on which case-control samples are available. While one may compare the significant variants of one trait with those of another, it is worth noting that the discovery of the variants in the first place may be under-powered in individual GWAS. Two other common strategies for examining genetic overlap between traits involve estimating genetic correlation, and testing how well polygenic risk score of one disease explains variation of the other. Both these approaches describe an overall genetic sharing, and do not indicate genetic sharing at a locus level or implicate novel shared variants or loci. Currently, there is no summary statistics based method to specifically test for pleiotropy between any two traits. Furthermore, there is no method for identifying pleiotropic loci between case-control traits that may or may not share controls.

In this article, we propose a formal statistical test of pleiotropy of two traits borrowing ideas from statistical mediation analysis literature. The proposed method, PLACO (pleiotropic analysis under composite null hypothesis), can be applied to summary-level data available from GWAS of two traits and can account for potential correlation across traits, such as that arising due to shared controls in case-control studies. We conduct extensive simulation experiments to study type I error and power of PLACO at stringent significance levels. We apply PLACO to summary data on common variants from two large case-control GWAS of European ancestry on Type 2 Diabetes (T2D) and on Prostate Cancer (PrCa). Many previous studies have reported an inverse association of these two chronic diseases suggesting shared risk factors; however, shared genetic mechanisms underlying this T2D-PrCa association is poorly understood. We replicate some candidate and known shared genes, and identify a number of novel shared genetic regions.

## Material and Methods

### Model and Notation

Consider two genome-wide studies of traits *Y*_1_ and *Y*_2_ on *n*_1_ and *n*_2_ individuals respectively who were genotyped and/or imputed or sequenced at *p* SNPs. Assume *n*_1_ individuals are independent of *n*_2_ individuals, with no overlapping samples between the studies. Let ***Y***_*k*_ and ***X***_*k*_ be the vectors of *k*-th trait values and genotypes at a given genetic variant respectively on all *n_k_* individuals (*k* = 1, 2). For the ease of explanation, we will assume the two traits are binary (e.g., case-control traits); however, our approach being based on summary statistics is applicable to two qualitative and/or quantitative traits. An individual’s outcome or trait can take value 0 for controls or 1 for cases. If the genetic variant is a bi-allelic single nucleotide polymorphism (SNP), an individual’s genotype can take value 0, 1 or 2 depending on the number of copies of minor alleles at the SNP. If the variant is imputed, the genotypic value will range between 0 and 2. For simplicity, we assume there is no covariate. Note, this assumption can be easily relaxed by considering trait residuals (obtained from regressing the covariates on the trait) instead of the raw trait values. Although residualizing outcome data is not standard, previous studies have shown that it does not affect validity of genetic association tests^26–28^.

The typical approach in a GWAS is to test for association of each genetic variant with the trait and report the estimated genetic effect sizes, their standard errors and the corresponding p-values for all genetic variants (often referred to as ‘summary statistics’). For a given genetic variant, the marginal model for outcome data is

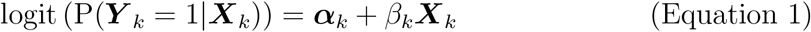

where *β_k_* is the genetic effect on the *k*-th trait (*k* = 1, 2). The null hypothesis of no association of the genetic variant with the *k*-th trait corresponds to 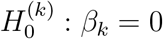. The Wald test statistic 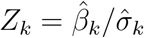 is used to test 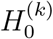, where 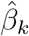 is the maximum likelihood estimate (MLE) of *β_k_* and 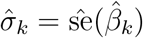 is its estimated standard error. For common variants, the *Z*-score (*Z_k_*) has an asymptotic *N*(0,1) distribution under the null 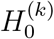. Since the two studies are assumed to be independent, *Z*_1_ and *Z*_2_ are expected to be independently distributed. It is to be noted that the Z-scores can also be obtained under any other genetic model (e.g., dominant or recessive), and the following methodological development is still applicable.

### Statistical Framework for Formal Testing of Pleiotropy

#### Defining the null hypothesis

The conventional cross-phenotype association methods test the global null hypothesis that none of the traits is associated with the given genetic variant (i.e., *β*_1_ = *β*_2_ = 0). Rejection of this global null can be due to one associated trait (*β*_1_ ≠ 0, *β*_2_ = 0 or *β*_1_ ≠ 0, *β*_2_ = 0). Here, we are interested in identifying the genetic variants that are associated with both the traits or outcomes (i.e., *β*_1_ ≠ 0, *β*_2_ ≠ 0). The effects of such a genetic variant on the traits may or may not be equal. Formally, our null hypothesis of no pleiotropy is *H*_0_: at most 1 trait is associated with the genetic variant while the alternative hypothesis is *H_a_*: both traits are associated.

#### A simple approach for testing pleiotropy

Mathematically, our null hypothesis of no pleiotropy is a composite null hypothesis *H*_0_: *H*_00_ ∪ *H*_01_ ∪ *H*_02_ while the alternative hypothesis is 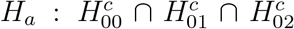, where *H*_00_: *β*_1_ = 0 = *β*_2_, *H*_01_: *β*_1_ = 0, *β*_2_ ≠ 0, *H*_02_: *β*_1_ = 0, *β*_2_ = 0 and 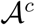 denotes the complement of set 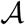. Thus, the alternative hypothesis is simply *H_a_*: *β*_1_ ≠ 0, *β*_2_ ≠ 0 (the situation we are interested in identifying). This is a special two-parameter case of the intersection-union principle of statistical hypothesis testing. A level-*α* intersection-union test (IUT)^29^ of *H*_0_ vs. *H_a_* is, reject *H*_0_ if a level-*α* test rejects *H*_0*k*_ for every *k* = 1,2. Consequently, the p-value for the IUT ≤ maximum of the p-values for testing 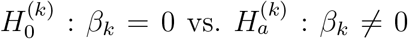. Thus, an approximate conservative p-value of the IUT is max {*p*_1_,*p*_2_}, where *p_k_* is the p-value corresponding to test statistic *Z_k_* (*k* = 1, 2). We refer to this approximate test as ‘maxP’ test in our figures and tables.

#### Other suitable approaches for testing pleiotropy

Observe that our null hypothesis of no pleiotropy can simply be written as *H*_0_: *β*_1_*β*_2_ = 0 vs. the alternative hypothesis *H_a_*: *β*_1_*β*_2_ ≠ 0. This immediately reminds us of the product of coefficients hypothesis tests for the significance of mediation effects in epidemiology^30^. It involves constructing test statistics by dividing 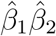 by its standard error and comparing the observed value of the test statistic to a standard normal distribution. Several variants of the standard error of 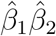 are used based on different assumptions and order of derivatives in the approximations. If Sobel’s approach^30,31^ is used in our context to test *H*_0_, the test statistic is 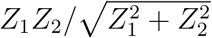, which uses an asymptotic *N*(0,1) distribution as its null distribution.

In the context of genome-wide mediation analysis, the normal approximation of Sobel’s method depends on a condition that only holds if at least one of the mediation coefficients is non-zero^32^. In the context of our pleiotropy test in GWAS, we expect most genetic variants to be not associated with either of the traits (i.e., we expect the global null *H*_00_ to be true for most genetic variants). As a consequence of sparse signals and hence the breakdown of condition for asymptotic normality of Sobel’s method, testing pleiotropy using Sobel’s method fails to control type I error and lacks power to detect pleiotropic effects of a genetic variant. In the mediation literature, as an alternative to Sobel’s method, Huang^32^ proposed a modified p-value calculation for the test of estimated mediation effect that maintains appropriate type I error under the assumption that most of the significance tests of mediation are conducted under the global null that both coefficients are zero. In this article, we borrow Huang’s approach^32^ from mediation analysis to propose a new single-variant test of pleiotropy of two traits. Our approach for identifying pleiotropic variants is particularly useful for characterizing genetic overlap between two disease traits from case-control GWAS at a variant level.

### Our Proposed Test of Pleiotropy: PLACO

#### Two independent traits

Suppose the global null *H*_00_ holds with probability *π*_00_ under which the single-trait test statistics *Z*_1_ and *Z*_2_ have asymptotic standard normal distributions. Further assume that the sub-null hypothesis *H*_01_ holds with probability *π*_01_ under which *Z*_1_ has a standard normal distribution and *Z*_2_ has a conditional *N*(*μ*_2_,1) distribution given the mean parameter *μ*_2_. We assume a 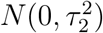 distribution for *μ*_2_. Similarly, assume that the sub-null hypothesis *H*_02_ holds with probability *π*_02_ and *Z*_2_ ~ *N*(0,1) while *Z*_1_|*μ*_1_ ∼ *N*(*μ*_1_,1), where 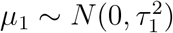.

In other words, we are assuming (a) *Z*_1_ and *Z*_2_ are independent *N*(0,1) variables under *H*_00_; (b) *Z*_1_ and *Z*_2_ are independent *N*(0,1) and 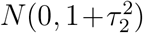 variables respectively under *H*_01_; and (c) *Z*_1_ and *Z*_2_ are independent 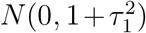 and *N*(0,1) variables respectively under *H*_02_. Consequently, the products 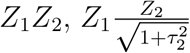 and 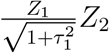 have normal product distributions under *H*_00_, *H*_01_ and *H*_02_ respectively (assuming the parameters *τ*_1_ and *τ*_2_ are known). The (symmetric) normal product distribution is given by the probability density function (p.d.f.) 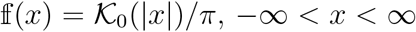, where 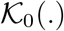 is the modified Bessel function of the second kind with order 0.

The p-value (two-tailed) for testing *H*_0_: *β*_1_*β*_2_ = 0 (no pleiotropy) against *H_a_*: *β*_1_*β*_2_ = 0 using the product of Z-scores as our test statistic is given by

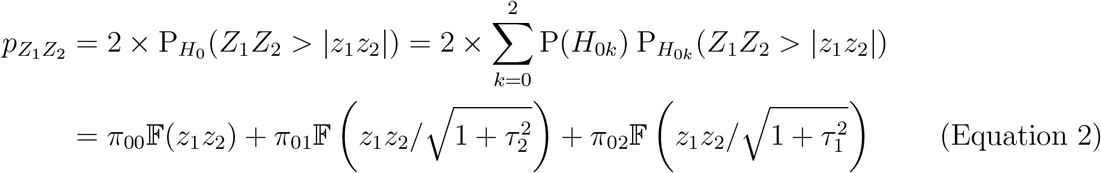

where *z*_1_ and *z*_2_ are the observed *Z*-scores for the two traits at a given genetic variant, and 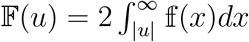 is the two-sided tail probability of a normal product distribution at value *u*. Observe that the analytical form for PLACO p-value (Equation 2) contains unknown parameters *π*_00_, *π*_01_, *π*_02_, *τ*_1_ and *τ*_2_. One can estimate these parameters only once under the null using the GWAS summary statistics on the millions of genetic variants genome-wide and assume they are known. However, this p-value evaluation approach is sensitive to these parameter estimates and can be quite conservative at genome-wide levels (Supplementary S1). Instead we will use an approximate asymptotic p-value to test the null hypothesis of no pleiotropy.

#### Asymptotic approximation of PLACO p-value

The PLACO p-value in Equation 2 can be approximated as

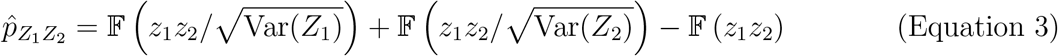

where 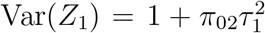 and similarly Var(*Z*_2_) are the estimated marginal variances of the *Z*-scores under the hierarchical model we assumed^32^. This can be implemented using our `R`^33^ program `PLACO` (https://github.com/RayDebashree/PLACO). The approximate *p*-value 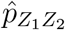 remains unchanged when mixture normal distributions or uniform distributions for the mean parameters μ_1_ and *μ_2_* (under *H*_02_ and *H*_01_ respectively) are assumed^32^.

#### Adjusting PLACO for correlation across GWAS

The above formulation of PLACO assumes that the *Z*-scores for the two traits are independent. While the independence of the effects 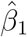 and 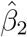, and consequently the Z-scores, is guaranteed in a mediation analysis^34^, it is not guaranteed for a pleiotropy analysis. If the two traits come from studies with overlapping samples, either partially (e.g. studies with shared controls^35,36^) or completely, then the *Z*-scores will be correlated^37^ and may lead to inflated p-values if the correlation is not accounted for in the pleiotropic analysis.

For two outcomes from two case-control studies, the correlation between the *Z*-scores is 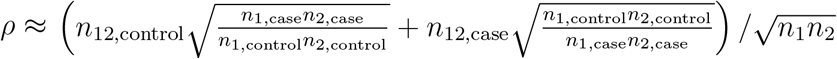, ignoring the variation due to 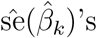, under the global null of no association, where *n*_*k*,case_ and *n*_*k*,control_ are respectively the number of cases and the number of controls in the study for *k*-th outcome, and *n*_12, control_ (*n*_12,case_) is the number of shared controls (cases) between the two studies^37^. In reality, the cases in two case-control studies are always independent and the control group in each study is at least as large as the case group. The correlation ρ, thus, ranges between 0 and 0.5, where the maximum is reached when there are equal number of cases and controls in each study, both studies have the same sample size and all the controls are shared (Supplementary S2). For two continuous traits, the correlation between the *Z*-scores is 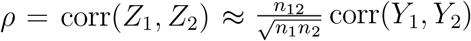 under the global null of no association, where *n*_12_ is the total number of overlapping samples (i.e., individuals with measurements on both traits) and *n*_1_, *n*_2_ are the respective sample sizes of the two traits^37^.

The number of overlapping samples between studies/traits may not be known when only GWAS summary data are available. In such a situation, one can estimate the correlation parameter *ρ* by the Pearson correlation of the *Z*-scores for the genetic variants with “no effect” on any trait. For a real dataset, the truth about which genetic variants have “no effect” is unknown. We choose the genetic variants that do not exceed a pre-defined significance threshold (say, genetic variants with single-trait p-value > 10^-4^) for any trait to estimate the correlation *ρ* between *Z*-scores^38^. One may also use cross-trait LD-score regression^39^ to estimate *ρ*; however we did not find appreciable differences between GWAS results obtained using estimates from these two approaches^13^. Irrespective of the approach, this estimation is done only once, as implemented in PLACO software, before applying PLACO genome-wide. If ***Z*** = (*Z*_1_, *Z*_2_) be the vector of Z-scores for a given genetic variant and 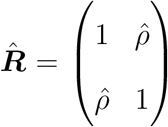 be the estimated correlation matrix, one needs to de-correlate the *Z*-scores as ***Z***^decor^ = ***R***^1/2^***Z*** so that 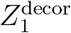 and 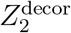 are uncorrelated. PLACO, as described before, can now be applied on these de-correlated *Z*-scores to test for pleiotropy of two correlated traits. However, we found from our simulation experiments that PLACO is an appropriate test of pleiotropy of two independent or moderately correlated traits, and may show inflated type I error for strongly correlated traits or when studies share more than half of their samples.

### Simulation Experiments

To evaluate operating characteristics of PLACO as a test for pleiotropy, we conduct simulation experiments in `R`^33^. We consider three broad simulation settings: one where we have traits from independent case-control studies, another with traits from case-control studies with shared controls, and the other with correlated traits from quantitative studies. For simplicity, we do not simulate any covariate or confounder. We simulate unrelated individuals and 10 million independent bi-allelic genetic variants in Hardy-Weinberg equilibrium with a fixed population-level minor allele frequency (MAF) 5%. We assume the commonly used additive genetic model in our simulations. Since we need multiple independent replicates to assess type I error control and power at stringent error thresholds, we assume the genetic variants are independent. Subsequently, we calculate estimated type I error (power) by averaging over the number of independent null (non-null) variants identified as having significant pleiotropic effect on both traits at a fixed significance level *α*.

Out of the 10 million genetic variants, we assume 99% variants to be under the global null of no association *H*_00_ (i.e., none of the two traits is associated with the genetic variant), 0.5% variants under the sub-null *H*_01_ (i.e., only the second trait is associated with the genetic variant), 0.4% variants under the sub-null *H*_02_ (i.e., only the first trait is associated with the genetic variant), and 0.1% variants under the alternative *H_a_* (i.e., the genetic variant has pleiotropic effect on both traits). Thus, our simulated dataset has 9.99 million null variants to estimate type I error and 10, 000 non-null variants to estimate statistical power.

#### Scenario I: Traits from two independent case-control studies

We simulate the two case-control studies such that the individuals in one study are independent of the other. We consider situations where the two studies have either comparable (1: 1) or unbalanced (4: 1) sample sizes. In other words, either the two studies have equal sample sizes (*n*_1_ = *n*_2_ = 2000) or the first study on the first trait is 4 times larger than the second study on the second trait (*n*_1_ = 8000, *n*_2_ = 2000). We assume a case-control ratio of 1: 1 in each study, and a baseline disease prevalence of 15% and 10% for the first and the second disease trait respectively. In this scenario, we compare type I error and power of Sobel’s approach, maxP approach, and PLACO to detect pleiotropy of the two independent case-control outcomes. The null genetic variants with non-zero effect on one trait only are assumed to have an odds ratio (OR) of 1.15 for the associated trait. For the non-null variants used to estimate power, we consider different choices of the two ORs to incorporate traits with genetic effects of varying directions and/or magnitudes.

#### Scenario II: Traits from two case-control studies with overlapping controls

We assume either 20%, 40%, 80% or 100% of the controls are shared, assuming equal number of controls in the two studies. Here, we compare type I error of Sobel’s approach, maxP approach, and PLACO with and without correction for sample overlap. Evaluating power in this scenario is redundant since the power will depend on the total number of independent samples, which we explore in Scenario I. For implementing PLACO that accounts for the overlap, we assume the number of overlapping samples is not available to calculate correlation through the Lin-Sullivan approach, and instead estimate the Pearson correlation of the *Z*-scores.

#### Scenario III: Two correlated traits from a study of quantitative traits

We simulate a single study with measurements on two correlated quantitative traits measured either on the same individuals (*n*_1_ = *n*_2_ = 2000) or the first trait is measured on many additional individuals (*n*_1_ = 8000, *n*_2_ = 2000). We vary both the strength and the direction of pairwise trait correlation: *ρ_trait_* = {–0.9, –0.4,0, 0.4, 0.9}. The null genetic variants with non-zero effect on one trait only are assumed to explain 0.1% of the variance of the associated trait. In this scenario, we only compare type I error of Sobel’s approach, maxP approach, and PLACO (with and without correction for correlation), and do not evaluate power.

### Application to T2D and Prostate Cancer GWAS Summary Data

Many epidemiologic studies^40–44^ of T2D and Prostate Cancer (PrCa) have reported association between these two diseases, suggesting shared risk factors. A few studies^45–48^ have been undertaken to identify shared genetic risk factors underlying this T2D-PrCa association. To elucidate shared genetic mechanisms between these two diseases, which is still poorly understood, we use our statistical approach PLACO on summary data from two of the largest and most recent GWAS of T2D and of PrCa in individuals of European ancestry.

Xue et al.^49^ meta-analyzed 62, 892 T2D cases and 596, 424 controls from three large GWAS datasets of European ancestry (DIAGRAM^50^, GERA^51^ and UK Biobank^52^). The authors reported summary statistics on 5,053,015 genotyped (from GWAS chip and Metabochip) and imputed autosomal SNPs (GRCh37/hg19) with MAF ⩾ 1% that were common to the three datasets. All imputed SNPs have imputation info score ⩾ 0.3. The reported summary statistics were obtained by fixed effects inverse-variance meta-analysis of GWAS summary statistics from each dataset after adjusting for study-specific covariates such as age, sex and principal components (PCs).

Schumacher et al.^53^ meta-analyzed 79,194 PrCa cases and 61,112 controls from eight GWAS or high-density SNP panels of European ancestry imputed to 1000 Genomes Phase 3. All imputed SNPs have imputation r^2^ ⩾ 0.3. The authors combined the per-allele odds ratios and standard errors, adjusted for PCs and study-relevant covariates, for the SNPs from the Illumina OncoArray and each GWAS by fixed effects inverse-variance meta-analysis. The summary statistics file contained information on 20, 370, 947 SNPs (GRCh37/hg19) across the autosomes and the X chromosome.

In this paper, we use the two sets of meta-analysis summary statistics of genetic association with T2D and with PrCa to detect shared common SNPs. We remove any SNP with allele mismatch between the two datasets, and focus on the remaining 5, 041, 948 autosomal SNPs with MAF ⩾ 1% that are common to both the datasets. For a given SNP, we harmonize the same effect allele across the two studies so that Z-scores from the two datasets can be jointly analyzed appropriately using PLACO. From the effect estimates and the standard errors, we calculate the *Z*-scores, and remove SNPs with *Z*^2^ > 80^54,55^ since extremely large effect sizes can disproportionately influence our analysis. The component studies underlying the T2D and the PrCa GWAS do not seem to overlap. The estimated correlation between *Z*-scores from T2D and those from PrCa is approximately 0 as well.

To characterize the findings from PLACO, we clump all the significantly associated SNPs (*ρ_PLACO_* < 5 × 10^-8^) in ±500 Kb radius and linkage disequilibrium (LD) threshold of *r*^2^ > 0.2 into a single genetic locus using `FUMA` (`SNP2GENE` function, v1.3.5e). We perform different gene-set enrichment analyses (using `GENE2FUNC` function) where the genes were prioritized by `FUMA` based on the loci identified by PLACO. To provide additional evidence of sharing at these loci, we perform Bayesian colocalization test^56^ of the PrCa and the T2D summary data using R package `coloc` (v3.2.1). This test computes 5 different overall posterior probabilities of the chosen region: 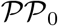 (posterior probability of no association with either disease), 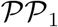 (association with T2D, not with PrCa), 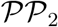 (association with PrCa, not with T2D), 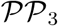 (association with both T2D and PrCa due to two distinct causal SNPs) and 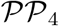 (association with both T2D and PrCa due to one common causal SNP). For each locus, we choose all the SNPs in ±200 Kb radius of the lead SNP and declare ‘convincing evidence’ of pleiotropic association of this region if it shows 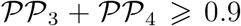 and 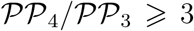 (cutoffs previously used elsewhere^57,58^). For this analysis, we use the `coloc.abf()` function on the effect estimates and their variance estimates for the SNPs in the chosen region (for each of T2D and PrCa) with default parameters and priors. For the significant loci with convincing evidence of colocalization, we manually look up Open Targets^59^ genetics platform to gather information about diseases associated with nearby genes (selected options ‘genetic associations’, ‘pathways & systems biology’ and ‘RNA expression’), and on relevant mouse data if available. To characterize the regulatory effects of the significant pleiotropic signals, we perform whole blood *cis* expression quantitative trait locus (eQTL) analysis using data from the eQTLGen Consortium^60^, the largest publicly available meta-analysis of blood eQTLs based on > 31, 500 individuals, in `FUMA`. For *cis*-eQTL analysis, we additionally consider T2D-relevant tissues (liver, pancreas, adipose, skeletal muscle)^61^ and PrCa-relevant tissue (prostate) from GTEx v8^62^.

## Results

### Simulation Experiments: Type I Error

#### Scenario I: Traits from two independent case-control studies

Irrespective of whether the sample sizes of the two studies are same or widely different, PLACO has well-calibrated type I error at stringent significance levels (Figure 1). In comparison, the other approaches are extremely conservative.

**Figure 1.**
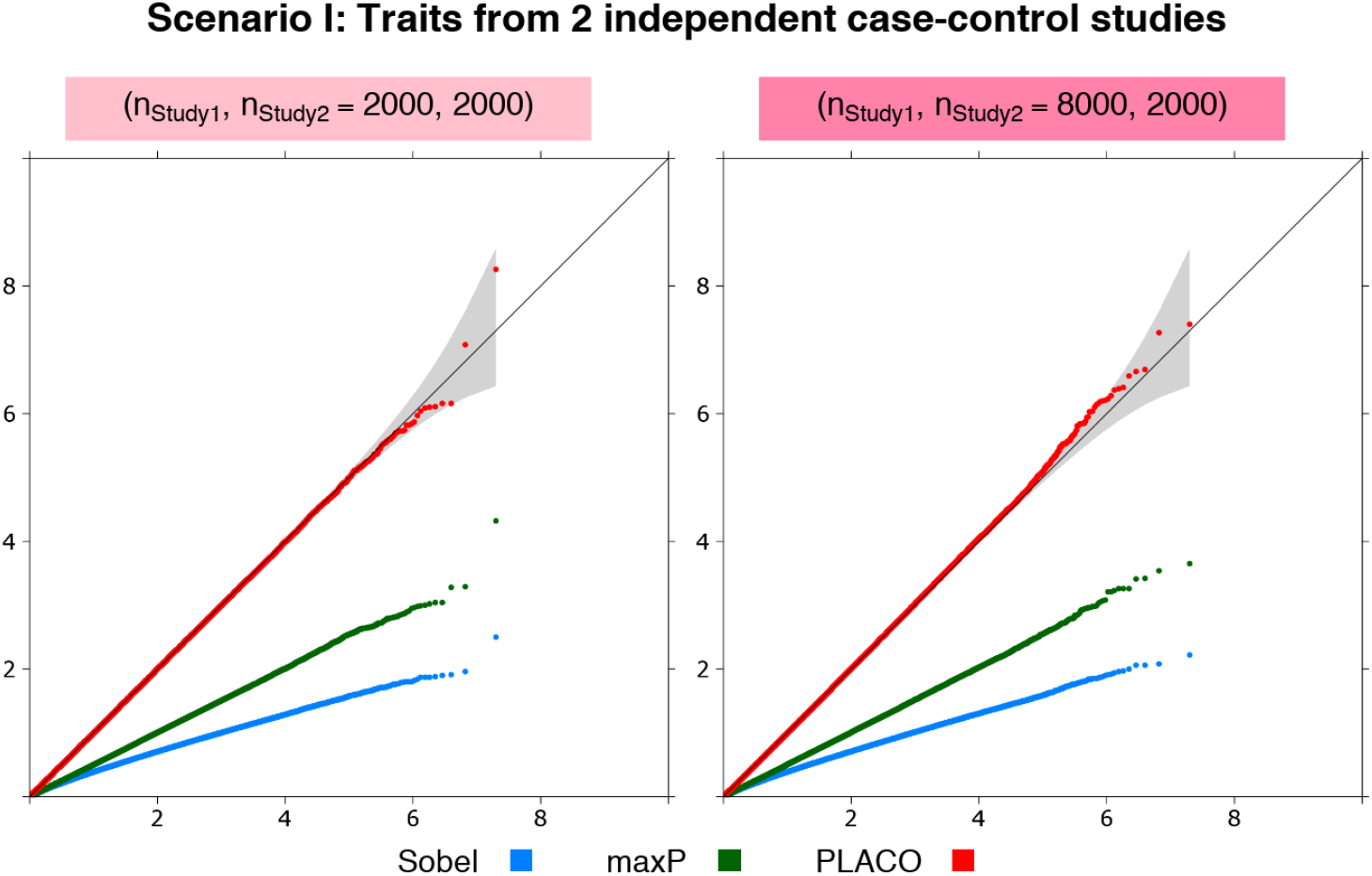
Scenario I: QQ plots for null data on traits from 2 independent case-control studies. Observed(- log_10_p-values) are plotted on the y-axis and Expected(- log_10_p-values) on the x-axis. Either each study has 1, 000 unrelated cases and 1,000 unrelated controls, or Study 1 is 4 times that of Study 2, where Study 2 has 1, 000 unrelated cases and 1, 000 unrelated controls. Type I error performance of tests of pleiotropic effect of a genetic variant on the 2 traits is based on 9.99 million null variants with genetic effects that are either {*β*_1_ = 0 = *β*_2_} or {*β*_1_ = 0,*β*_2_ = log(1.15)} or {*β*_1_ = log(1.15),*β*_2_ = 0}. The gray shaded region represents a conservative 95% confidence interval for the expected distribution of p-values. P-values ⩾ 10^-10^ are shown here.

#### Scenario II: Traits from two case-control studies with overlapping controls

Regardless of the extent of control overlap in the two studies, PLACO exhibits appropriate type I error when correlation is accounted for in the analysis (Figures 2 and S2). We also note that if *Z*-scores are not decorrelated for studies with overlapping samples, pleiotropy analysis will likely show spurious association signals as indicated by the inflated ‘PLACO (no overlap correction)’ curve. The other approaches are still very conservative across all scenarios of overlap.

**Figure 2.**
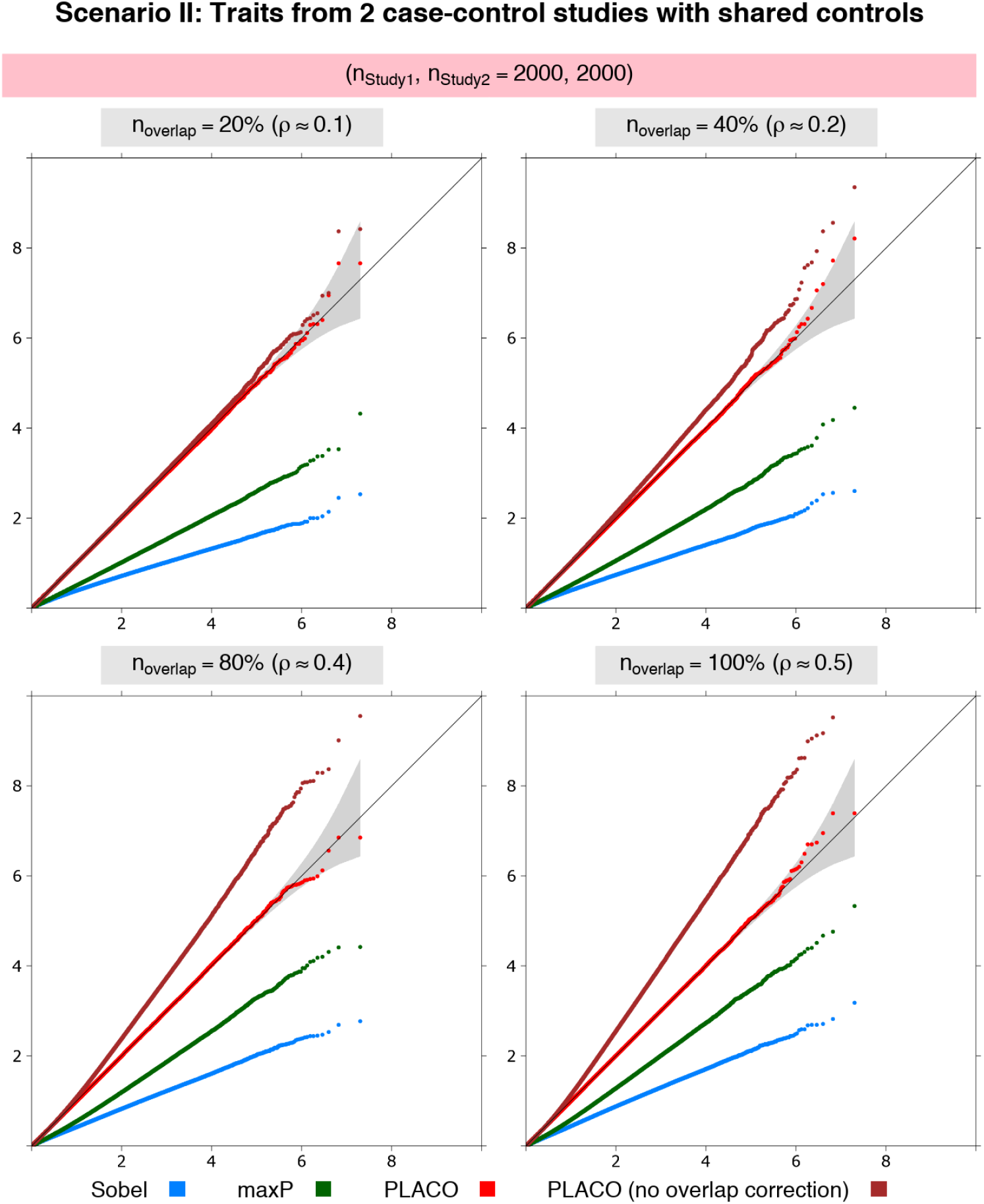
Scenario II: QQ plots for null data on traits from 2 case-control studies with different proportions of overlapping controls. Observed(– log_10_p-values) are plotted on the y-axis and Expected(– log_10_p-values) on the x-axis. Equal study sample size, and equal casecontrol size assumed in each study. Each study has 1,000 unrelated cases and 1, 000 unrelated controls, of which either 20%, 40%, 80% or 100% of the controls are shared between the two studies. Type I error performance of tests of pleiotropic effect of a genetic variant on the 2 traits is based on 9.99 million null variants with genetic effects that are either {*β*_1_ = 0 = *β*_2_} or {*β*_1_ = 0, *β*_2_ = log(1.15)} or {*β*_1_ = log(1.15), *β*_2_ = 0}. The gray shaded region represents a conservative 95% confidence interval for the expected distribution of p-values. P-values ⩾ 10^-10^ are shown here.

#### Scenario III: Two correlated traits from a study of quantitative traits

We find PLACO has well-calibrated type I error for moderately correlated traits irrespective of the direction of correlation between the traits, and has inflated type I error for strongly correlated traits (Figure S3). Application of PLACO ignoring correlation will show spurious association signals. As before, the other approaches exhibit conservative behavior across all scenarios of pairwise trait correlation. The ‘maxP’ approach can, however, be less conservative for strongly correlated traits.

### Simulation Experiments: Power

#### Scenario I: Traits from two independent case-control studies

For benchmarking, we compare power of PLACO against the naive approach of declaring pleiotropy when a variant reaches genome-wide significance for the first trait with the larger sample size and reaches a more liberal significance threshold for the second trait. We use two such naive approaches: one using criterion *p*_Trait1_ < 5 × 10^-8^, *p*_Trait2_ < 10^-5^ and the other *p*_Trait1_ < 5 × 10^-8^, *p*_Trait2_ < 10^-3^ (‘Naive-1’ and ‘Naive-2’ respectively in our figures). We do not compare with any of the other methods because they do not maintain appropriate type I error. Regardless of the magnitude and directions of pleiotropic association and the sample size differences between studies, PLACO has dramatically improved statistical power to detect pleiotropy compared to the naive approaches (Figure 3). When testing pleiotropy in a real dataset, all the other approaches are expected to lack power due to their very conservative type I error control.

**Figure 3.**
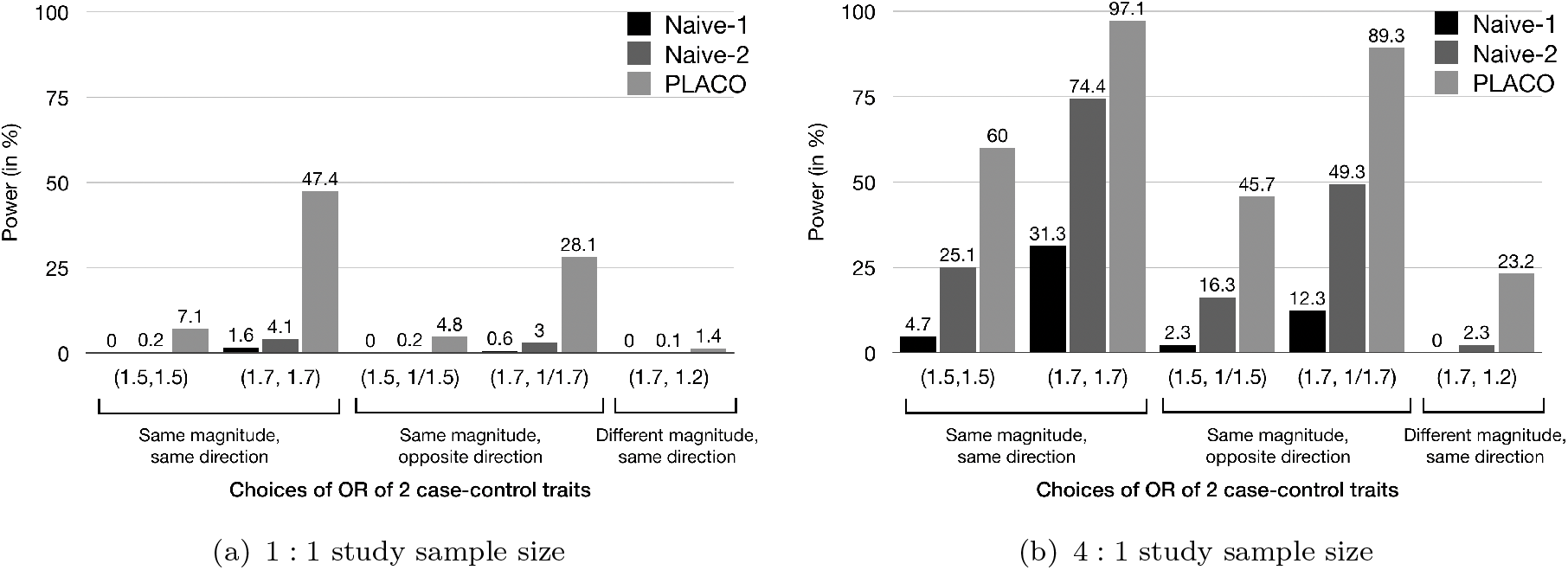
Scenario I: Power of PLACO and naive approaches at genome-wide significance level (5 × 10^-8^) for varying genetic effects of traits from 2 independent case-control studies. The first naive approach (‘Naive-1’) declares pleiotropic association when *p*_Trait1_ < 5 × 10^-8^ and *p*_Trait2_ < 10^-5^, while the second naive approach (‘Naive-2’) uses a more liberal criterion *p*_Trait1_ < 5 × 10^-8^ and *p*_Trait2_ < 10^-3^. (a) Each study either has 1, 000 unrelated cases and 1, 000 unrelated controls, or (b) Study 1 has 4 times sample size as Study 2, where Study 2 has 1, 000 unrelated cases and 1, 000 unrelated controls.

### Application to T2D and Prostate Cancer GWAS Summary Data

#### Overview of joint T2D-PrCa locus level associations

PLACO identified 1, 329 genomewide significant SNPs that mapped to 44 distinct loci (Figure 4). The lead SNPs of 24 loci (55%) increase risk for one outcome while decreasing risk for the other. This observation is consistent with what observational studies^41,63,64^ and genetic risk-score studies^46,47^ have reported before: an inverse association between T2D and PrCa. We define a locus as novel if there is no ‘previously associated SNP’ from GWAS catalog^65^ (as of December 16, 2019) within ±500 Kb radius or in LD (r^2^ > 0.2) with our index SNP, the GWAS peak, from that locus. To define ‘previously associated SNP’ in our context of pleiotropy of T2D and PrCa, we looked for any SNP within each locus that is associated with both T2D-related trait (either of T2D, 2-hour glucose challenge, glucose level, glycated albumin, HbA1c, insulin level, proinsulin level, insulin resistance, insulin response, or glycemic traits) and PrCa-related trait (either of PrCa or prostate-specific antigen levels). Since GWAS catalog includes exome-wide studies, we chose a slightly liberal exome-wide significance threshold of p < 5 × 10^-7^ to define association. We discovered 38 potentially novel loci, after liftover of GRCh38 genomic coordinates in GWAS catalog to hg19 using R package `liftOver`^66^.

**Figure 4.**
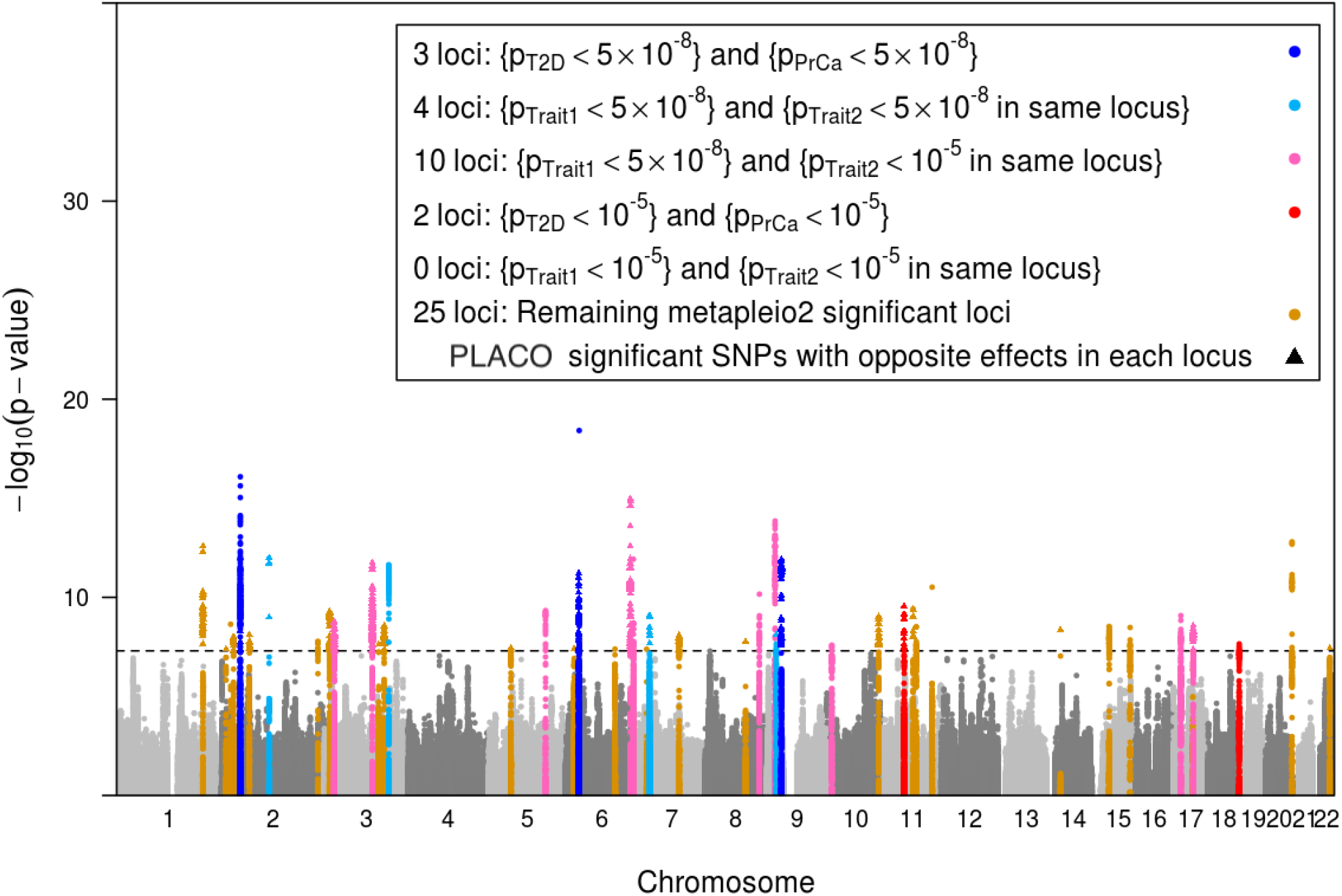
Manhattan Plot of the PLACO p-values of pleiotropic association of common genetic variants with outcomes (traits) T2D and PrCa. The black horizontal dashed line corresponds to genome-wide significance level *α* = 5 × 10^-8^. The 44 loci with genome-wide significant pleiotropic lead SNP have been highlighted. A locus is defined by clumping SNPs in ±500 Kb radius around the lead significant SNP and with LD *r*^2^ > 0.2. Within each locus, if a PLACO significant SNP has genetic effects in opposite directions for T2D and PrCa, it is plotted as a solid triangle (24 such loci), else as a solid circle. Each identified pleiotropic locus is categorized (color-coded) as follows. Three loci harbor SNPs that are genome-wide significant for both T2D and PrCa (single-trait *p* < 5 × 10^-8^). Four loci contain SNPs that are genome-wide significant for one disease and in close proximity (i.e., in the same locus) with another SNP genome-wide significant for the other disease. There are 10 loci where SNPs are genome-wide significant for one disease and in close proximity with another SNP suggestively significant (single-trait *p* < 10^-5^) for the other disease. Two loci harbor SNPs that are suggestively significant (but not genome-wide significant) for both T2D and PrCa. There is no locus that contain SNPs that are suggestively significant (but not genome-wide significant) for one disease and in close proximity with another SNP suggestively significant (but not genome-wide significant) for the other disease. The rest of the 25 loci identified by PLACO contain SNPs that are not even suggestively significant for either T2D or PrCa.

#### PLACO points to known and candidate shared genetic regions

GWAS catalog search reveals that 6 out of 44 loci near genes *THADA, BCL2L11, AC005355.2, PBX2* (in the major histo-compatibility complex or MHC region of 6p21), *JAZF1* and *CDKN2A/B* have been previously implicated in studies of both T2D and PrCa. In particular, *THADA*^43^ (Figure S6) and *JAZF1* ^45^ (Figure S7) represent well-recognized shared genetic regions between T2D and PrCa. *HNF1B,* also known as *TCF2,* is another recognized shared gene^45,67^, which we fail to detect possibly because we excluded SNPs with extremely large effect sizes^54,55^ (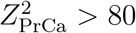 for many SNPs positionally mapped in/near *HNF1B*), which may have weakened any signal in this region. Signals from PLACO point to candidate shared genes such as *PPARG*^47^ (Figure S8) and *CDKN2A*^43,47^ (Figure S9). PLACO did not find enough evidence of shared genetic component in other previously explored genes such as *KCNQ1* ^43^ (Figure S10) and *MTNR1B*^43^ (Figure S11).

#### Gene-set enrichment analysis

For further analysis, we exclude the 1 locus that lay in the MHC region of chromosome 6p21 because of strong SNP associations in this long-range and complex LD block that complicates fine-mapping efforts^61^. The 310 genes to which the 43 pleiotropic loci were mapped by `FUMA` are significantly enriched in GWAS catalog reported genes for PrCa, T2D and other T2D related traits (Figure S12). When tested for tissue specificity against differentially expressed genes from GTEx v8 data across 53 tissue types, these genes are significantly enriched in pancreas (a T2D-relevant tissue) and whole-blood (Figure S13). Analyses in other annotated gene sets from Molecular Signatures Database (MSigDB v7.0)^68^ and in curated biological pathways from WikiPathways^69^, and functional enrichment analyses are described in Supplementary S5.

#### Colocalization analysis

Bayesian colocalization tests of ±200 Kb region around the lead SNPs of the 43 loci reveal 26 lead SNPs as having the highest posterior probability of being associated with both PrCa and T2D (Table 1). Eight loci show convincing evidence of containing SNPs that are likely causal for both T2D and PrCa, 7 of which have the highest posterior probabilities of being causal SNPs and exhibit stronger signals of pleiotropic association than the single trait associations (Table 2). The lead SNP for the eighth locus, near *RGS17,* is 54 Kb away from the SNP with the highest causal probability (rs6932847), and both have similar PLACO p-value of pleiotropic association.

**Table 1.**
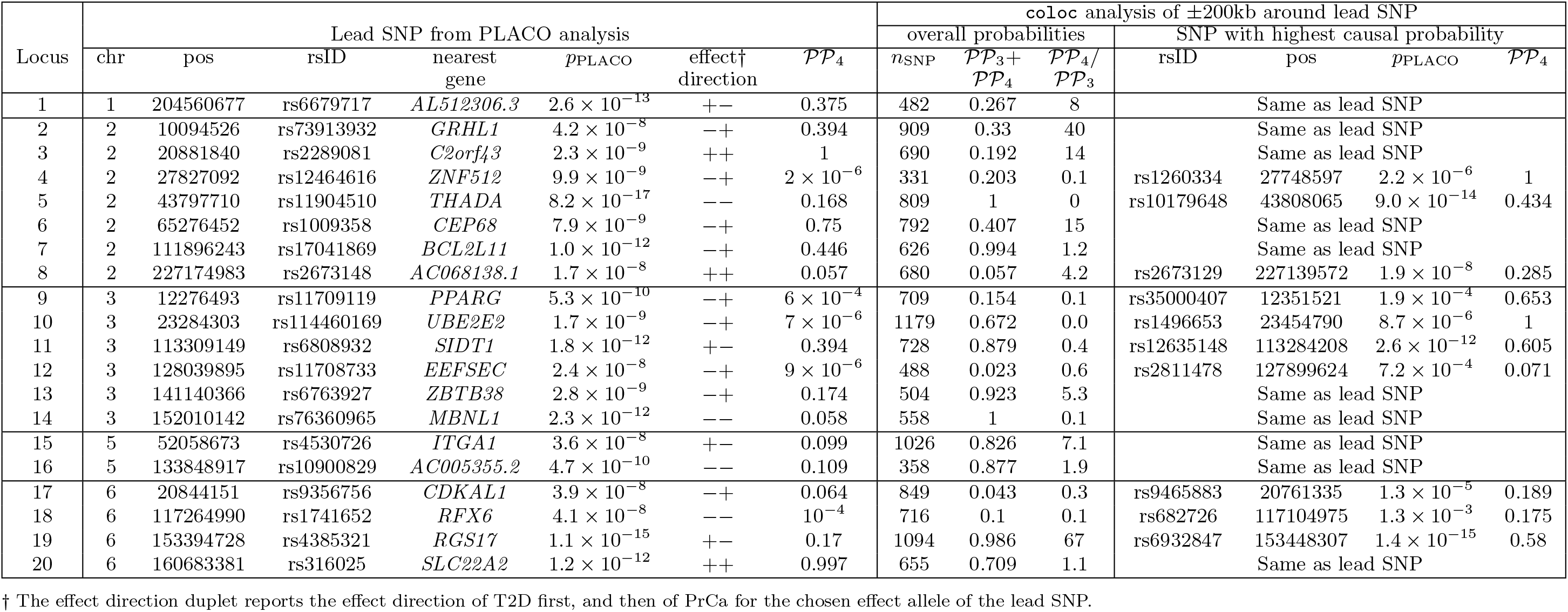
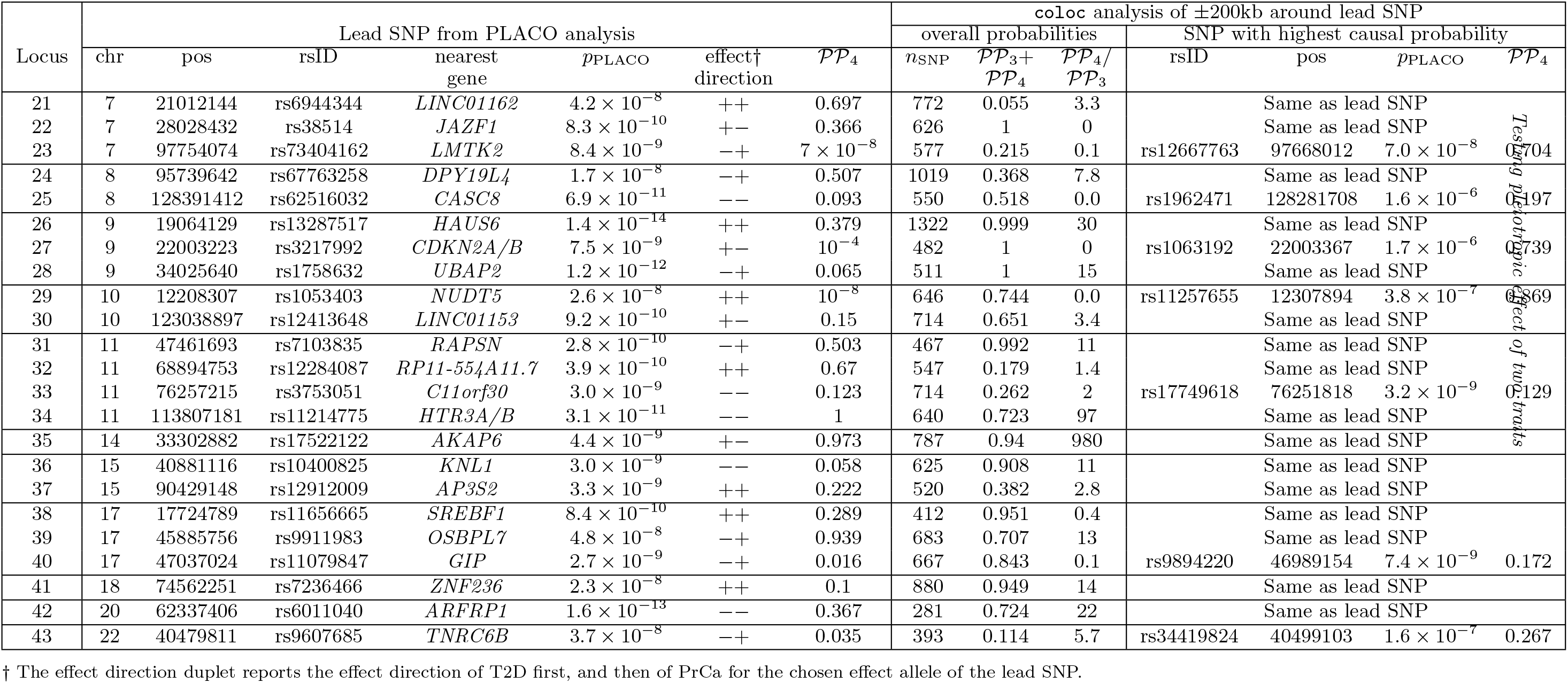
*This table lists the coloc colocalization posterior probability 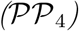 for the lead SNPs from each of the* 43 *loci identified by PLACO. A high 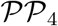 for a SNP indicates high probability of being causal for both T2D and PrCa. SNPs with highest 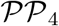 within ±200 Kb of the lead SNPs are also reported.*

**Table 2.** *This table lists the potentially novel loci detected by PLACO and with convincing evidence* (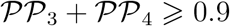 *and* 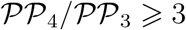) *of being causal for both T2D and PrCa from colocalization analysis*.

| Genomic Locus | Lead SNP from PLACO analysis |  |  |  |  |  |  | Summary statistics for lead SNP |  |  |  |  |  |
| --- | --- | --- | --- | --- | --- | --- | --- | --- | --- | --- | --- | --- | --- |
| | chr | pos | rsID | nearest gene | effect allele | other allele | effect allele freq. | CADD score | $\beta_{T2D}$ | $p_{T2D}$ | $\beta_{PrCa}$ | $p_{PrCa}$ | $p_{PLACO}$ |
| 13 | 3 | 141140366 | rs6763927 | ZBTB38 | T | A | 0.44 | 3.18 | -0.0316 | $6.8 \times 10^{-5}$ | 0.0459 | $8.5 \times 10^{-9}$ | $2.8 \times 10^{-9}$ |
| 19 | 6 | 153394728 | rs4385321 | RGS17 | A | G | 0.35 | 4.05 | 0.0352 | $2.8 \times 10^{-6}$ | -0.0724 | $2.7 \times 10^{-18}$ | $1.1 \times 10^{-15}$ |
| 26 | 9 | 19064129 | rs13287517 | HAUS6 | C | G | 0.39 | 0.44 | 0.0402 | $5.3 \times 10^{-7}$ | 0.0609 | $7.1 \times 10^{-14}$ | $1.4 \times 10^{-14}$ |
| 28 | 9 | 34025640 | rs1758632 | UBAP2 | C | G | 0.38 | 1.24 | -0.0491 | $1.4 \times 10^{-9}$ | 0.0432 | $1.1 \times 10^{-7}$ | $1.2 \times 10^{-12}$ |
| 31 | 11 | 47461693 | rs7103835 | RAPSN | A | G | 0.31 | 7.53 | -0.0384 | $1.2 \times 10^{-6}$ | 0.046 | $1.4 \times 10^{-7}$ | $2.9 \times 10^{-10}$ |
| 35 | 14 | 33302882 | rs17522122 | AKAP6 | T | G | 0.48 | 2.19 | 0.0403 | $5.2 \times 10^{-8}$ | -0.0337 | $4.0 \times 10^{-5}$ | $4.4 \times 10^{-9}$ |
| 36 | 15 | 40881116 | rs10400825 | KNL1 | G | A | 0.15 | 2.66 | -0.0452 | $4.0 \times 10^{-5}$ | -0.0612 | $2.4 \times 10^{-8}$ | $3.0 \times 10^{-9}$ |
| 41 | 18 | 74562251 | rs7236466 | ZNF236 | G | T | 0.38 | 4.03 | 0.0368 | $3.8 \times 10^{-6}$ | 0.0364 | $8.2 \times 10^{-6}$ | $2.3 \times 10^{-8}$ |

#### Characterizing the 8 most interesting potentially novel pleiotropic loci

The lead SNPs of 6 of the 8 potentially novel pleiotropic loci with convincing evidence from the colocalization analyses have effect alleles that increase risk for one disease while protecting from the other (Table 2). While the 8 loci contain *cis*-eQTLs in multiple T2D-relevant tissues (Figures S17-S22), SNPs in the loci near *RGS17* (Figure 5) and *UBAP2* (Figure 6) show significant *cis*-eQTL associations in both T2D-relevant and PrCa-relevant tissues. In Open Targets, genes near the *ZBTB38, UBAP2* and *ZNF236* loci show associations with various cancers, diabetes and obesity (no relevant mouse data available for these genes). The *RGS17* locus show associations with various cancers, including PrCa and prostate neoplasm, and body mass index (BMI) but has no known associations with any T2D-related trait (no relevant mouse data available). Of particular interest are the *HAUS6* and the *RAPSN* loci. While *HAUS6* and its nearby genes *RRAGA* and *PLIN2* have various cancers (including PrCa) as associated diseases in Open Targets, one or more of them are related to metabolism phenotype, abnormal gluconeogenesis and hypoglycemia in mice. GWAS catalog search of these genes did not yield any known association result with any T2D-related trait. Similarly, the nearby gene *MADD* for the *RAPSN* locus has various cancers, neoplasms and glucose-related phenotypes as associated diseases in Open Targets; and is a recognized T2D gene, which when knocked out in mice, show impaired glucose tolerance, hyperglycemia and abnormal pancreatic beta cell morphology.

**Figure 5.**
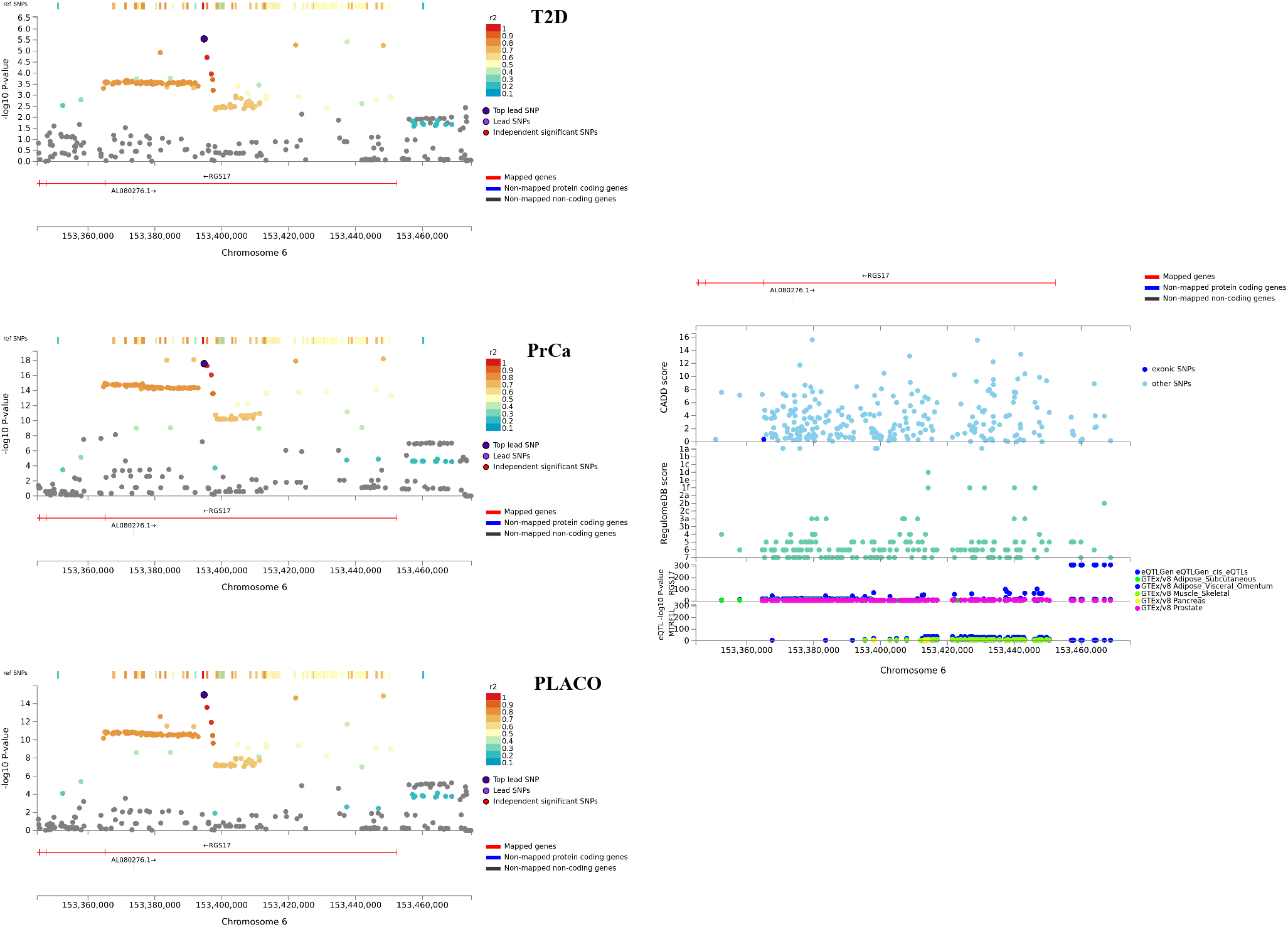
Regional association plot of significant pleiotropic locus near *RGS17* with annotations such as CADD scores, RegulomeDB scores, and *cis* eQTL association p-values from 6 tissues (whole blood from eQTLGen Consortium; and adipose, liver, muscle-skeletal, pancreas, and prostate tissues from GTEx v8.

**Figure 6.**
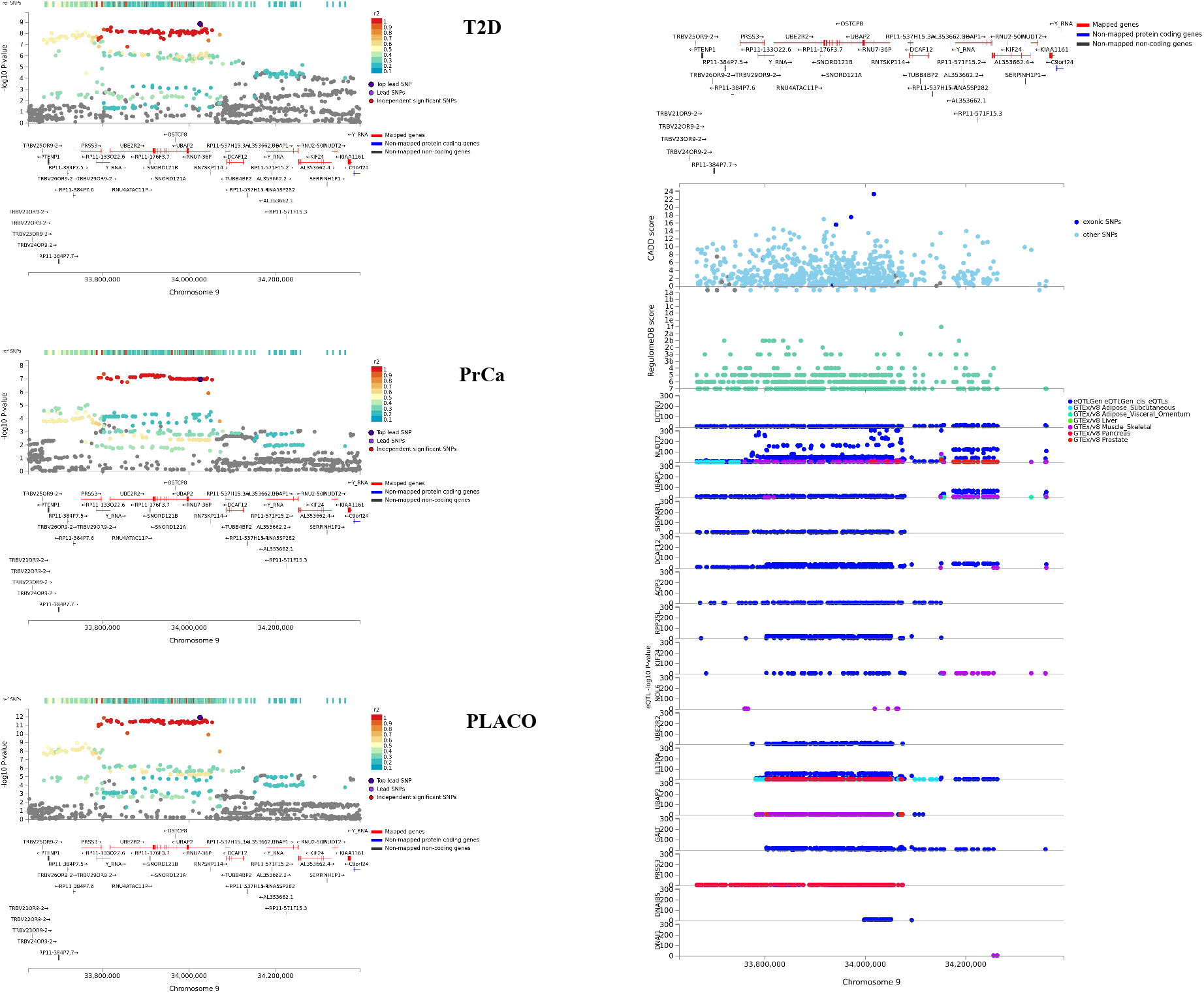
Regional association plot of significant pleiotropic locus near *UBAP2* with annotations such as CADD scores, RegulomeDB scores, and *cis* eQTL association p-values from 6 tissues (whole blood from eQTLGen Consortium; and adipose, liver, muscle-skeletal, pancreas, and prostate tissues from GTEx v8.

## Discussion

In this paper, we propose a formal statistical hypothesis test and a novel method, PLACO, to determine common pleiotropic or shared variants of two independent traits and show how it may well be applied to correlated traits or traits from studies with sample overlap. In our simulations involving qualitative and quantitative traits with unequal prevalences, unequal genetic effect sizes, unequal sample sizes – ranging from modest to large – and with/without overlapping samples, PLACO exhibits well-calibrated type I error. We find PLACO is powerful in detecting subtle genetic effects of pleiotropic variants that may or may not be in the same direction and that may be missed when each disease trait is analyzed separately (some additional simulations in Supplementary S4). Statistical power is significantly improved when PLACO is used, compared to the naive approach that identifies pleiotropy when a genetic variant reaches genome-wide significance for the trait with larger sample size and reaches a more liberal threshold for the other. Based on our simulations, we advocate using PLACO on independent traits, or moderately correlated traits after decorrelating the Z-scores as described before.

We use the most recent publicly available case-control GWAS summary data on T2D and on PrCa in individuals of European ancestry to determine variants that influence risk to both these diseases. We identify several known and candidate shared genes, and detect a number of novel shared genetic regions near *ZBTB38* (3q23), *RGS17* (6q25.3), *HAUS6* (9p22.1), *UBAP2* (9p13.3), *RAPSN* (11p11.2), *AKAP6* (14q12), *KNL1* (15q15) and *ZNF236* (18q23). A recent study^70^ showed a weak positive genetic correlation between T2D and PrCa. It is worth noting that the concept of genetic correlation is different from pleiotropy. For genetic correlation to be non-zero, the directions of effect of non-null variants must be consistently aligned^39^. Effect alleles of at least half of the significant SNPs identified by PLACO have opposite genetic effects on the two diseases, which supports many previous studies reporting inverse relationship between T2D and PrCa, and likely explains the weak genetic correlation in the previous study.

The key advantage of PLACO is not requiring individual-level data which makes it easily applicable to datasets for which only GWAS summary data are available. It does not require compute intensive permutations or Monte Carlo simulations to calculate p-value of simultaneous association of two traits with one genetic variant. We are conveniently using the asymptotic normality of MLE of genetic effects to get at the null distribution of the PLACO test statistic. The existence of an analytical form for PLACO p-value (Equation 2) and its approximation (Equation 3) makes it suitable for application on a genome-wide scale. While we have applied PLACO to summary statistics from population-based case-control GWAS, it may also be applied to summary statistics of two traits from family-based designs (e.g., disease traits from case-parent trio studies). For instance, family-based GWAS data from several study cohorts will soon be available under the cohort collaboration study, Environmental influences on Child Health Outcomes (ECHO, https://www.nih.gov/research-training/ environmental-influences-child-health-outcomes-echo-program), to understand genetic underpinnings of pediatric outcomes. One important scientific question will be to identify genetic overlap of such outcomes (e.g., neurodevelopmental disorders, respiratory disorders), which PLACO can conveniently address that too without having to pool individual-level data.

Our study and our statistical approach are not without limitations. PLACO requires genome-wide summary data to infer pleiotropic association of each variant, and cannot be used when summary data on only a handful of candidate genetic variants are available. PLACO shows inflated type I error when the traits are strongly correlated even after using our decorrelation approach. The approximate PLACO p-value 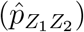 is a good approximation when the non-zero effect under *H*_01_ or *H*_02_ is small^32^, else it may be inflated. Simply stated, if the effect of a genetic variant is very strong on one trait and has no effect on the other trait, the p-value reported by `PLACO` may be inflated and indicate a genome-wide significant result. We suggest that SNPs with marginal *Z*^2^ > 80 be removed before analysis, similar to suggestion for LD-score regression approaches. PLACO is a single-variant association test that is not expected to control type I error for genetic variants with low minor allele counts since the asymptotic normality of MLE assumption may be violated. It can only detect statistical association of a variant with two traits, referred to as ‘statistical pleiotropy’^58^, and cannot distinguish between the various types of pleiotropy: biological, mediated, spurious due to design artefacts or spurious due to strong LD between causal variants in different genes ^1^. Notwithstanding these caveats, PLACO provides massive power gain over commonly used approaches, and shows promise in providing additional evidence for a shared genetic component between two traits.

## Supplemental Data

Supplemental Data include additional text, figures and tables, and can be found with this article online.

## Acknowledgments

This research was supported by the NIH for the Environmental influences of Child Health Outcomes (ECHO) Data Analysis Center (U24OD023382). It was carried out in part using computing cluster – the Joint High Performance Computing Exchange (JHPCE) – at the Department of Biostatistics, Johns Hopkins Bloomberg School of Public Health. Part of this research project was conducted using computational resources at the Maryland Advanced Research Computing Center (MARCC).

## Declaration of Interests

The authors declare no competing interest.

## Web Resources

PLACO, https://github.com/RayDebashree/PLACO (coming soon)

Type 2 diabetes summary data, http://cnsgenomics.com/data/t2d

Prostate cancer summary data, ftp://ftp.ebi.ac.uk/pub/databases/gwas/summary_statistics/SchumacherFR_29892016_GCST006085

Bayesian colocalization analysis, https://cran.r-project.org/web/packages/coloc FUMA, https://fuma.ctglab.nl/

QQ plot code, https://genome.sph.umich.edu/wiki/Code_Sample:_Generating_QQ_Plots_in_R

Manhattan plot code, https://genome.sph.umich.edu/wiki/Code_Sample:_Generating_Manhattan_Plots_in_R

Locuszoom plot, http://locuszoom.org/

## Notes

### Competing Interest Statement

The authors have declared no competing interest.

## References

1. Solovieff, N., Cotsapas, C., Lee, P. H., Purcell, S. M., and Smoller, J. W. Pleiotropy in complex traits: challenges and strategies. Nat Rev Genet, 14(7):483–495, 2013.

2. Sivakumaran, S., Agakov, F., Theodoratou, E., Prendergast, J. G., Zgaga, L., Manolio, T., Rudan, I., McKeigue, P., Wilson, J. F., and Campbell, H. Abundant pleiotropy in human complex diseases and traits. Am J Hum Genet, 89(5):607–618, 2011.

3. Wu, Y.-H., Graff, R. E., Passarelli, M. N., Hoffman, J. D., Ziv, E., Hoffmann, T. J., and Witte, J. S. Identification of pleiotropic cancer susceptibility variants from genome-wide association studies reveals functional characteristics. Cancer Epidemiol Biomarkers Prev, 27(1):75–85, 2018.

4. Cotsapas, C., Voight, B. F., Rossin, E., Lage, K., Neale, B. M., Wallace, C., Abecasis, G. R., Barrett, J. C., Behrens, T., Cho, J., et al. Pervasive sharing of genetic effects in autoimmune disease. PLoS Genet, 7(8):e1002254, 2011.

5. Cross-Disorder Group of the Psychiatric Genomics Consortium. Identification of risk loci with shared effects on five major psychiatric disorders: a genome-wide analysis. Lancet, 381(9875):1371–1379, 2013.

6. Amare, A. T., Vaez, A., Hsu, Y.-H., Direk, N., Kamali, Z., Howard, D. M., McIntosh, A. M., Tiemeier, H., Bültmann, U., Snieder, H., et al. Bivariate genome-wide association analyses of the broad depression phenotype combined with major depressive disorder, bipolar disorder or schizophrenia reveal eight novel genetic loci for depression. Mol Psychiatry, page 1, 2019.

7. Li, R., Brockschmidt, F. F., Kiefer, A. K., Stefansson, H., Nyholt, D. R., Song, K., Vermeulen, S. H., Kanoni, S., Glass, D., Medland, S. E., et al. Six novel susceptibility loci for early-onset androgenetic alopecia and their unexpected association with common diseases. PLoS Genet, 8(5):e1002746, 2012.

8. Hui, K. Y., Fernandez-Hernandez, H., Hu, J., Schaffner, A., Pankratz, N., Hsu, N.-Y., Chuang, L.-S., Carmi, S., Villaverde, N., Li, X., et al. Functional variants in the *LRRK2* gene confer shared effects on risk for Crohn’s disease and Parkinson’s disease. Sci Transl Med, 10(423):eaai7795, 2018.

9. Pickrell, J. K., Berisa, T., Liu, J. Z., Ségurel, L., Tung, J. Y., and Hinds, D. A. Detection and interpretation of shared genetic influences on 42 human traits. Nat Genet, 48(7): 709–717, 2016.

10. Yang, C., Li, C., Wang, Q., Chung, D., and Zhao, H. Implications of pleiotropy: challenges and opportunities for mining Big Data in biomedicine. Front Genet, 6:229, 2015.

11. Gratten, J. and Visscher, P. M. Genetic pleiotropy in complex traits and diseases: implications for genomic medicine. Genome Med, 8(1):78, 2016.

12. Hackinger, S. and Zeggini, E. Statistical methods to detect pleiotropy in human complex traits. Open Biol, 7(11):170125, 2017.

13. Ray, D. and Chatterjee, N. Effect of non-normality and low count variants on cross phenotype association tests in GWAS. Eur J Hum Genet, 28:300–312, 2020.

14. Baker, A. R., Goodloe, R. J., Larkin, E. K., Baechle, D. J., Song, Y. E., Phillips, L. S., and Gray-McGuire, C. L. Multivariate association analysis of the components of metabolic syndrome from the Framingham Heart Study. In BMC Proc, volume 3, page S42. BioMed Central, 2009.

15. Inouye, M., Ripatti, S., Kettunen, J., Lyytikäinen, L.-P., Oksala, N., Laurila, P.-P., Kangas, A. J., Soininen, P., Savolainen, M. J., Viikari, J., et al. Novel loci for metabolic networks and multi-tissue expression studies reveal genes for atherosclerosis. PLoS Genet, 8(8):e1002907, 2012.

16. Medina-Gomez, C., Kemp, J. P., Dimou, N. L., Kreiner, E., Chesi, A., Zemel, B. S., Bønnelykke, K., Boer, C. G., Ahluwalia, T. S., Bisgaard, H., et al. Bivariate genome-wide association meta-analysis of pediatric musculoskeletal traits reveals pleiotropic effects at the *SREBF1/TOM1L2* locus. Nat Commun, 8(1):121, 2017.

17. Heid, I. M. and Winkler, T. W. A multitrait GWAS sheds light on insulin resistance. Nat Genet, 49(1):7, 2017.

18. Shen, X., Klarić, L., Sharapov, S., Mangino, M., Ning, Z., Wu, D., Trbojević-Akmačić, I., Pučić-Baković, M., Rudan, I., Polašek, O., et al. Multivariate discovery and replication of five novel loci associated with immunoglobulin GN-glycosylation. Nat Commun, 8(1): 447, 2017.

19. Zhao, W., Rasheed, A., Tikkanen, E., Lee, J.-J., Butterworth, A. S., Howson, J. M., Assimes, T. L., Chowdhury, R., Orho-Melander, M., Damrauer, S., et al. Identification of new susceptibility loci for type 2 diabetes and shared etiological pathways with coronary heart disease. Nat Genet, 49(10):1450–1457, 2017.

20. Baselmans, B. M., Jansen, R., Ip, H. F., van Dongen, J., Abdellaoui, A., van de Weijer, M. P., Bao, Y., Smart, M., Kumari, M., Willemsen, G., et al. Multivariate genome-wide analyses of the well-being spectrum. Nat Genet, 51(3):445–451, 2019.

21. Nath, A. P., Ritchie, S. C., Grinberg, N. F., Tang, H. H., Huang, Q. Q., Teo, S. M., Ahola-Olli, A. V., Würtz, P., Havulinna, A. S., Aalto, K., et al. Multivariate genomewide association analysis of a cytokine network reveals variants with widespread immune, haematological, and cardiometabolic pleiotropy. Am J Hum Genet, 105(6):1076–1090, 2019.

22. Zhang, Q., Feitosa, M., and Borecki, I. B. Estimating and testing pleiotropy of single genetic variant for two quantitative traits. Genet Epidemiol, 38(6):523–530, 2014.

23. Schaid, D. J., Tong, X., Larrabee, B., Kennedy, R. B., Poland, G. A., and Sinnwell, J. P. Statistical methods for testing genetic pleiotropy. Genetics, 204(2):483–497, 2016.

24. Lutz, S. M., Fingerlin, T. E., Hokanson, J. E., and Lange, C. A general approach to testing for pleiotropy with rare and common variants. Genet Epidemiol, 41(2):163–170, 2017.

25. Schaid, D. J., Tong, X., Batzler, A., Sinnwell, J. P., Qing, J., and Biernacka, J. M. Multivariate generalized linear model for genetic pleiotropy. Biostatistics, 20(1):111–128, 2017.

26. Price, A. L., Patterson, N. J., Plenge, R. M., Weinblatt, M. E., Shadick, N. A., and Reich, D. Principal components analysis corrects for stratification in genome-wide association studies. Nat Genet, 38(8):904–909, 2006.

27. Kang, H. M., Sul, J. H., Service, S. K., Zaitlen, N. A., Kong, S.-y., Freimer, N. B., Sabatti, C., Eskin, E., et al. Variance component model to account for sample structure in genome-wide association studies. Nat Genet, 42(4):348–354, 2010.

28. Broadaway, K. A., Cutler, D. J., Duncan, R., Moore, J. L., Ware, E. B., Jhun, M. A., Bielak, L. F., Zhao, W., Smith, J. A., Peyser, P. A., et al. A statistical approach for testing cross-phenotype effects of rare variants. Am J Hum Genet, 98(3):525–540, 2016.

29. Berger, R. L. Likelihood Ratio Tests and Intersection-Union Tests, pages 225–237. Birkhäauser Boston, Boston, MA, 1997.

30. MacKinnon, D. P., Lockwood, C. M., Hoffman, J. M., West, S. G., and Sheets, V. A comparison of methods to test mediation and other intervening variable effects. Psychol Methods, 7(1):83–104, 2002.

31. Sobel, M. E. Asymptotic confidence intervals for indirect effects in structural equation models. Sociol Methodol, 13:290–312, 1982.

32. Huang, Y.-T. Genome-wide analyses of sparse mediation effects under composite null hypotheses. Ann Appl Stat, 13(1):60–84, 2019.

33. R Core Team. R: A Language and Environment for Statistical Computing. R Foundation for Statistical Computing, Vienna, Austria, 2018. URL https://www.R-project.org/.

34. MacKinnon, D. P., Warsi, G., and Dwyer, J. H. A simulation study of mediated effect measures. Multivariate Behav Res, 30(1):41–62, 1995.

35. Wellcome Trust Case Control Consortium et al. Genome-wide association study of 14,000 cases of seven common diseases and 3,000 shared controls. Nature, 447(7145):661–678, 2007.

36. Mitchell, B. D., Fornage, M., McArdle, P. F., Cheng, Y.-C., Pulit, S., Wong, Q., Dave, T., Williams, S. R., Corriveau, R., Gwinn, K., et al. Using previously genotyped controls in genome-wide association studies (GWAS): application to the Stroke Genetics Network (SiGN). Front Genet, 5:95, 2014.

37. Lin, D.-Y. and Sullivan, P. Meta-analysis of genome-wide association studies with overlapping subjects. Am J Hum Genet, 85(6):862–872, 2009.

38. Ray, D. and Boehnke, M. Methods for meta-analysis of multiple traits using GWAS summary statistics. Genet Epidemiol, 42(2):134–145, 2018.

39. Bulik-Sullivan, B., Finucane, H. K., Anttila, V., Gusev, A., Day, F. R., Loh, P.-R., Consortium, R., Consortium, P. G., for Anorexia Nervosa of the Wellcome Trust Case Control Consortium 3, G. C., Duncan, L., Perry, J. R. B., Patterson, N., Robinson, E. B., Daly, M. J., Price, A. L., and Neale, B. M. An atlas of genetic correlations across human diseases and traits. Nat Genet, 47(11):1236–1241, 2015.

40. Giovannucci, E., Rimm, E. B., Stampfer, M. J., Colditz, G. A., and Willett, W. C. Diabetes mellitus and risk of prostate cancer (United States). Cancer Causes Control, 9(1):3–9, 1998.

41. Kasper, J. S. and Giovannucci, E. A meta-analysis of diabetes mellitus and the risk of prostate cancer. Cancer Epidemiol Biomarkers Prev, 15(11):2056–2062, 2006.

42. Waters, K. M., Henderson, B. E., Stram, D. O., Wan, P., Kolonel, L. N., and Haiman, C. A. Association of diabetes with prostate cancer risk in the multiethnic cohort. Am J Epidemiol, 169(8):937–945, 2009.

43. Machiela, M. J., Lindström, S., Allen, N. E., Haiman, C. A., Albanes, D., Barricarte, A., Berndt, S. I., Bueno-de Mesquita, H. B., Chanock, S., Gaziano, J. M., et al. Association of type 2 diabetes susceptibility variants with advanced prostate cancer risk in the Breast and Prostate Cancer Cohort Consortium. Am J Epidemiol, 176(12):1121–1129, 2012.

44. Gallagher, E. J. and LeRoith, D. Epidemiology and molecular mechanisms tying obesity, diabetes, and the metabolic syndrome with cancer. Diabetes Care, 36(Supplement 2): S233–S239, 2013.

45. Frayling, T., Colhoun, H., and Florez, J. A genetic link between type 2 diabetes and prostate cancer. Diabetologia, 51(10):1757–1760, 2008.

46. Pierce, B. L. and Ahsan, H. Genetic susceptibility to type 2 diabetes is associated with reduced prostate cancer risk. Hum Hered, 69(3):193–201, 2010.

47. Meyer, T. E., Boerwinkle, E., Morrison, A. C., Volcik, K. A., Sanderson, M., Coker, A. L., Pankow, J. S., and Folsom, A. R. Diabetes genes and prostate cancer in the Atherosclerosis Risk in Communities study. Cancer Epidemiol Biomarkers Prev, 19(2): 558–565, 2010.

48. Yu, O. H. Y., Foulkes, W. D., Dastani, Z., Martin, R. M., Eeles, R., Richards, J. B., Consortium, P., Investigators, C. G., et al. An assessment of the shared allelic architecture between type II diabetes and prostate cancer. Cancer Epidemiol Biomarkers Prev, 22(8):1473–1475, 2013.

49. Xue, A., Wu, Y., Zhu, Z., Zhang, F., Kemper, K. E., Zheng, Z., Yengo, L., Lloyd-Jones, L. R., Sidorenko, J., Wu, Y., et al. Genome-wide association analyses identify 143 risk variants and putative regulatory mechanisms for type 2 diabetes. Nat Commun, 9(1): 2941, 2018.

50. Morris, A. P., Voight, B. F., Teslovich, T. M., Ferreira, T., Segre, A. V., Steinthorsdottir, V., Strawbridge, R. J., Khan, H., Grallert, H., Mahajan, A., et al. Large-scale association analysis provides insights into the genetic architecture and pathophysiology of type 2 diabetes. Nat Genet, 44(9):981–990, 2012.

51. Banda, Y., Kvale, M. N., Hoffmann, T. J., Hesselson, S. E., Ranatunga, D., Tang, H., Sabatti, C., Croen, L. A., Dispensa, B. P., Henderson, M., et al. Characterizing race/ethnicity and genetic ancestry for 100,000 subjects in the Genetic Epidemiology Research on Adult Health and Aging (GERA) cohort. Genetics, 200(4):1285–1295, 2015.

52. Bycroft, C., Freeman, C., Petkova, D., Band, G., Elliott, L. T., Sharp, K., Motyer, A., Vukcevic, D., Delaneau, O., O’Connell, J., et al. The UK Biobank resource with deep phenotyping and genomic data. Nature, 562(7726):203–209, 2018.

53. Schumacher, F. R., Al Olama, A. A., Berndt, S. I., Benlloch, S., Ahmed, M., Saunders, E. J., Dadaev, T., Leongamornlert, D., Anokian, E., Cieza-Borrella, C., et al. Association analyses of more than 140,000 men identify 63 new prostate cancer susceptibility loci. Nat Genet, 50(7):928, 2018.

54. Bulik-Sullivan, B. K., Loh, P.-R., Finucane, H. K., Ripke, S., Yang, J., Patterson, N., Daly, M. J., Price, A. L., Neale, B. M., of the Psychiatric Genomics Consortium, S. W. G., et al. LD score regression distinguishes confounding from polygenicity in genome-wide association studies. Nat Genet, 47(3):291–295, 2015.

55. Zheng, J., Erzurumluoglu, A. M., Elsworth, B. L., Kemp, J. P., Howe, L., Haycock, P. C., Hemani, G., Tansey, K., Laurin, C., Early Genetics and Lifecourse Epidemiology (EAGLE) Eczema Consortium, et al. LD Hub: a centralized database and web interface to perform LD score regression that maximizes the potential of summary level GWAS data for SNP heritability and genetic correlation analysis. Bioinformatics, 33(2):272–279, 2016.

56. Giambartolomei, C., Vukcevic, D., Schadt, E. E., Franke, L., Hingorani, A. D., Wallace, C., and Plagnol, V. Bayesian test for colocalisation between pairs of genetic association studies using summary statistics. PLoS Genet, 10(5):e1004383, 2014.

57. Guo, H., Fortune, M. D., Burren, O. S., Schofield, E., Todd, J. A., and Wallace, C. Integration of disease association and eQTL data using a Bayesian colocalisation approach highlights six candidate causal genes in immune-mediated diseases. Hum Mol Genet, 24(12):3305–3313, 2015.

58. Watanabe, K., Stringer, S., Frei, O., Mirkov, M. U., de Leeuw, C., Polderman, T. J., van der Sluis, S., Andreassen, O. A., Neale, B. M., and Posthuma, D. A global overview of pleiotropy and genetic architecture in complex traits. Nat Genet, 51(9):1339–1348, 2019.

59. Koscielny, G., An, P., Carvalho-Silva, D., Cham, J. A., Fumis, L., Gasparyan, R., Hasan, S., Karamanis, N., Maguire, M., Papa, E., et al. Open Targets: a platform for therapeutic target identification and validation. Nucleic Acids Res, 45(D1):D985–D994, 2016.

60. Võsa, U., Claringbould, A., Westra, H.-J., Bonder, M. J., Deelen, P., Zeng, B., Kirsten, H., Saha, A., Kreuzhuber, R., Kasela, S., et al. Unraveling the polygenic architecture of complex traits using blood eQTL metaanalysis. bioRxiv, 2018.. URL https://www.biorxiv.org/content/early/2018/10/19/447367.

61. Mahajan, A., Taliun, D., Thurner, M., Robertson, N. R., Torres, J. M., Rayner, N. W., Payne, A. J., Steinthorsdottir, V., Scott, R. A., Grarup, N., et al. Fine-mapping type 2 diabetes loci to single-variant resolution using high-density imputation and islet-specific epigenome maps. Nat Genet, 50:1505–1513, 2018.

62. GTEx Consortium et al. Genetic effects on gene expression across human tissues. Nature, 550(7675):204–213, 2017.

63. Bonovas, S., Filioussi, K., and Tsantes, A. Diabetes mellitus and risk of prostate cancer: a meta-analysis. Diabetologia, 47(6):1071–1078, 2004.

64. Tande, A. J., Platz, E. A., and Folsom, A. R. The metabolic syndrome is associated with reduced risk of prostate cancer. Am J Epidemiol, 164(11):1094–1102, 2006.

65. Buniello, A., MacArthur, J. A. L., Cerezo, M., Harris, L. W., Hayhurst, J., Malangone, C., McMahon, A., Morales, J., Mountjoy, E., Sollis, E., et al. The NHGRI-EBI GWAS Catalog of published genome-wide association studies, targeted arrays and summary statistics 2019. Nucleic Acids Res, 47(D1):D1005–D1012, 2019.

66. Bioconductor Package Maintainer. liftOver: Changing genomic coordinate systems with rtracklayer::liftOver., 2019. URL https://www.bioconductor.org/help/workflows/liftOver/. R package version 1.10.0.

67. Gudmundsson, J., Sulem, P., Steinthorsdottir, V., Bergthorsson, J. T., Thorleifsson, G., Manolescu, A., Rafnar, T., Gudbjartsson, D., Agnarsson, B. A., Baker, A., et al. Two variants on chromosome 17 confer prostate cancer risk, and the one in TCF2 protects against type 2 diabetes. Nat Genet, 39(8):977–983, 2007.

68. Liberzon, A., Subramanian, A., Pinchback, R., Thorvaldsdóttir, H., Tamayo, P., and Mesirov, J. P. Molecular signatures database (MSigDB) 3.0. Bioinformatics, 27(12): 1739–1740, 2011.

69. Kutmon, M., Riutta, A., Nunes, N., Hanspers, K., Willighagen, E. L., Bohler, A., Mélius, J., Waagmeester, A., Sinha, S. R., Miller, R., et al. WikiPathways: capturing the full diversity of pathway knowledge. Nucleic Acids Res, 44(D1):D488–D494, 2015.

70. Lindstróm, S., Finucane, H., Bulik-Sullivan, B., Schumacher, F. R., Amos, C. I., Hung, R. J., Rand, K., Gruber, S. B., Conti, D., Permuth, J. B., et al. Quantifying the genetic correlation between multiple cancer types. Cancer Epidemiol Biomarkers Prev, 26(9): 1427–1435, 2017.

