## Supplemental Data for "A Powerful Method for Pleiotropic Analysis under Composite Null Hypothesis Identifies Novel Shared Loci Between Type 2 Diabetes and Prostate Cancer"

FOR

BY


Debashree Ray[1,2], Nilanjan Chatterjee[2,3]

[1]*Department of Epidemiology, and* [2]*Department of Biostatistics*

*Bloomberg School of Public Health, Johns Hopkins University, Baltimore, MD, USA.*

[3]*Department of Oncology, School of Medicine, Johns Hopkins University, Baltimore, MD, USA.*

#### Supplementary S1

**Estimation of PLACO p-value.** The analytical form for PLACO p-value in Equation 2 contains unknown parameters $\pi_{00}$, $\pi_{01}$, $\pi_{02}$, $\tau_1$ and $\tau_2$. To estimate these parameters, one may employ the approach taken by cross-phenotype summary-statistics based methods to estimate the covariance matrix of multiple $Z$-scores[1]. For instance, the proportion of genetic variants with marginal p-values $> 10^{-4}$ for both traits can be used as an estimate of $\pi_{00}$. An estimate of $\pi_{01}$ is the proportion of genetic variants with p-values $> 10^{-4}$ for the first trait and with p-values $< 10^{-4}$ for the second trait. $\pi_{02}$ can be similarly estimated. For estimating $\tau_1$, observe that the variance of $Z_1$ under $H_{02}$ is $1 + \tau_1^2$. Therefore, the estimated variance of $Z_1$ corresponding to genetic variants with p-values $< 10^{-4}$ for the first trait and with p-values $> 10^{-4}$ for the second trait can be used as an estimate of $1 + \tau_1^2$ in Equation 2. Similarly, the variance parameter $\tau_2$ can be estimated. Note that this estimation procedure (denoted as 'PLACO (estimated pvalue)' in the subsequent figures) is done only once for a given study using the single-trait $Z$-scores and p-values ('summary statistics') that are usually publicly available from a GWAS.

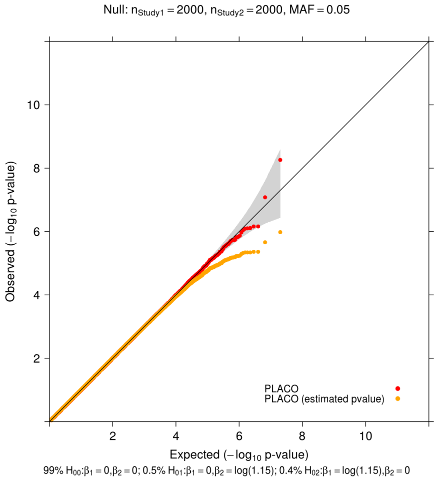


(a) 1 : 1 study sample size

**Figure S1:** Scenario I: Comparison of the PLACO approach using the asymptotic approximate p-value vs using the estimated p-value. QQ plots for null data are plotted on 2 case-control traits from 2 independent studies with fixed genetic effects. Observed($-\log_{10}$p-values) are plotted on the y-axis and Expected($-\log_{10}$p-values) on the x-axis. Each study has $1,000$ unrelated cases and $1,000$ unrelated controls. Performance of the tests of pleiotropic effect of a genetic variant on the 2 traits is based on 9.99 million variants. The gray shaded region represents a conservative 95% confidence interval for the expected distribution of p-values. P-values $\geq 10^{-12}$ are shown here.

#### Supplementary S2

**Correlation between case-control studies with shared controls.** For two outcomes from two case-control studies, the correlation between the $Z$-scores is

$$\rho \approx \left( n_{12,\text{co}} \sqrt{\frac{n_{1,\text{ca}} n_{2,\text{ca}}}{n_{1,\text{co}} n_{2,\text{co}}}} + n_{12,\text{ca}} \sqrt{\frac{n_{1,\text{co}} n_{2,\text{co}}}{n_{1,\text{ca}} n_{2,\text{ca}}}} \right) / \sqrt{n_1 n_2}$$

(ignoring the variation due to $\hat{\text{se}}(\hat{\beta}_k)$'s) under the global null of no association, where $n_{k,\text{ca}}$ and $n_{k,\text{co}}$ are respectively the number of cases and the number of controls in the study for $k$-th outcome, and $n_{12,\text{co}}$ ($n_{12,\text{ca}}$) is the number of shared controls (cases) between the two studies[2]. In reality, the cases in two case-control studies are always independent and the control group in each study is at least as large as the case group. Based on this, let us assume (1) 100% control overlap and no shared case ($n_{12,\text{ca}} = 0$); (2) the case:control ratio in both studies is the same (say, $1 : r_c$, where $r_c \geq 1$ - a reasonable assumption because the number of controls is almost always larger than the number of cases); (3) the total sample size of study 1 is $r_s$ times that of study 2, where $r_s \geq 1$. From assumption (1), the correlation boils down to $\rho \approx \left( n_{12,\text{co}} \sqrt{\frac{n_{1,\text{ca}} n_{2,\text{ca}}}{n_{1,\text{co}} n_{2,\text{co}}}} \right) / \sqrt{n_1 n_2}$. From assumptions (1) and (2), we get $n_{12,\text{co}} = n_{1,\text{co}} = n_{2,\text{co}}$ and $n_{j,\text{co}} = r_c n_{j,\text{ca}}$ for study $j = 1, 2$. From assumptions (2) and (3), we have $n_1 = r_s n_2$, $n_{1,\text{ca}} = r_s n_{2,\text{ca}}$, and $n_{1,\text{co}} = r_s n_{2,\text{co}}$. Finally, under these 3 assumptions, we have $\rho \approx \frac{1}{\sqrt{r_s}(1+r_c)}$, the maximum of which is attained when $r_s$ and $r_c$ take the lowest possible value. Thus, the correlation $\rho$ reaches a maximum of 0.5 when there are equal number of cases and controls in each study, both studies have the same sample size and all the controls (and no case) are shared.

**Table S1:** Possible values of correlation $\rho$ between $Z$-scores of two outcomes from two case-control studies with complete control overlap, with one study being $r_s(\geq 1)$ times as large as the other study, and with $r_c(\geq 1)$ controls for each case in both studies.

| $1 : r_c$ \ $r_s : 1$ | 1 : 1 | 1.5 : 1 | 2 : 1 | 3 : 1 | 5 : 1 | 10 : 1 |
|---|---|---|---|---|---|---|
| 1 : 1 | 0.5 | 0.41 | 0.35 | 0.29 | 0.22 | 0.16 |
| 1 : 4 | 0.2 | 0.16 | 0.14 | 0.11 | 0.09 | 0.06 |
| 1 : 9 | 0.1 | 0.08 | 0.07 | 0.06 | 0.04 | 0.03 |

### Supplementary S3

#### Additional figures and tables from simulation experiments.

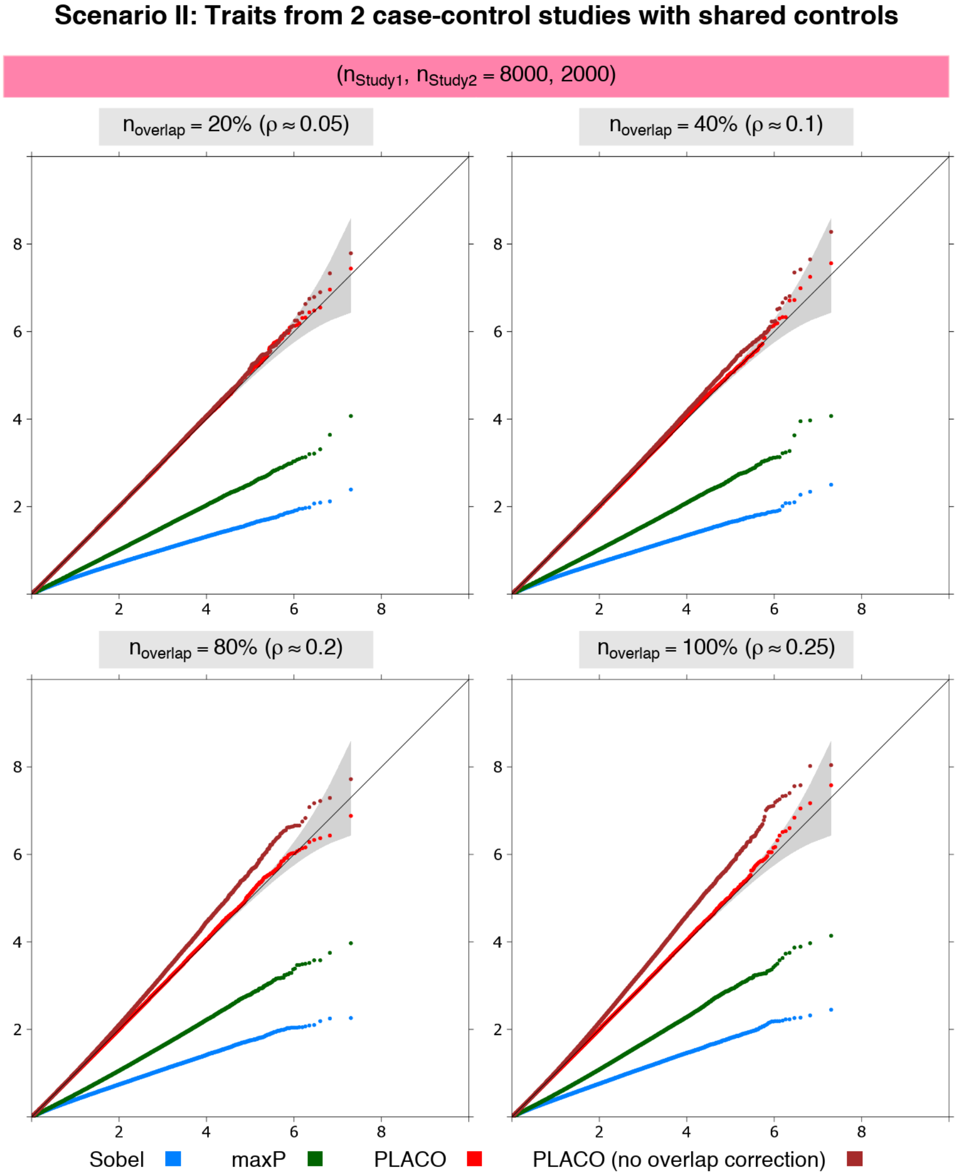


**Figure S2:** Scenario II: QQ plots for null data on traits from 2 case-control studies with different proportions of overlapping controls. Observed($-\log_{10}$p-values) are plotted on the y-axis and Expected($-\log_{10}$p-values) on the x-axis. Unequal study sample size, and equal case-control size assumed in each study. Study 1 has 4,000 unrelated cases and 4,000 unrelated controls. Study 2 has 1,000 unrelated cases and 1,000 unrelated controls, of which either 20%, 40%, 80% or 100% of the controls are shared between the two studies. Type I error performance of tests of pleiotropic effect of a genetic variant on the 2 traits is based on 9.99 million null variants with genetic effects that are either $\{\beta_1 = 0 = \beta_2\}$ or $\{\beta_1 = 0, \beta_2 = \log(1.15)\}$ or $\{\beta_1 = \log(1.15), \beta_2 = 0\}$. The gray shaded region represents a conservative 95% confidence interval for the expected distribution of p-values. P-values $\geq 10^{-10}$ are shown here.

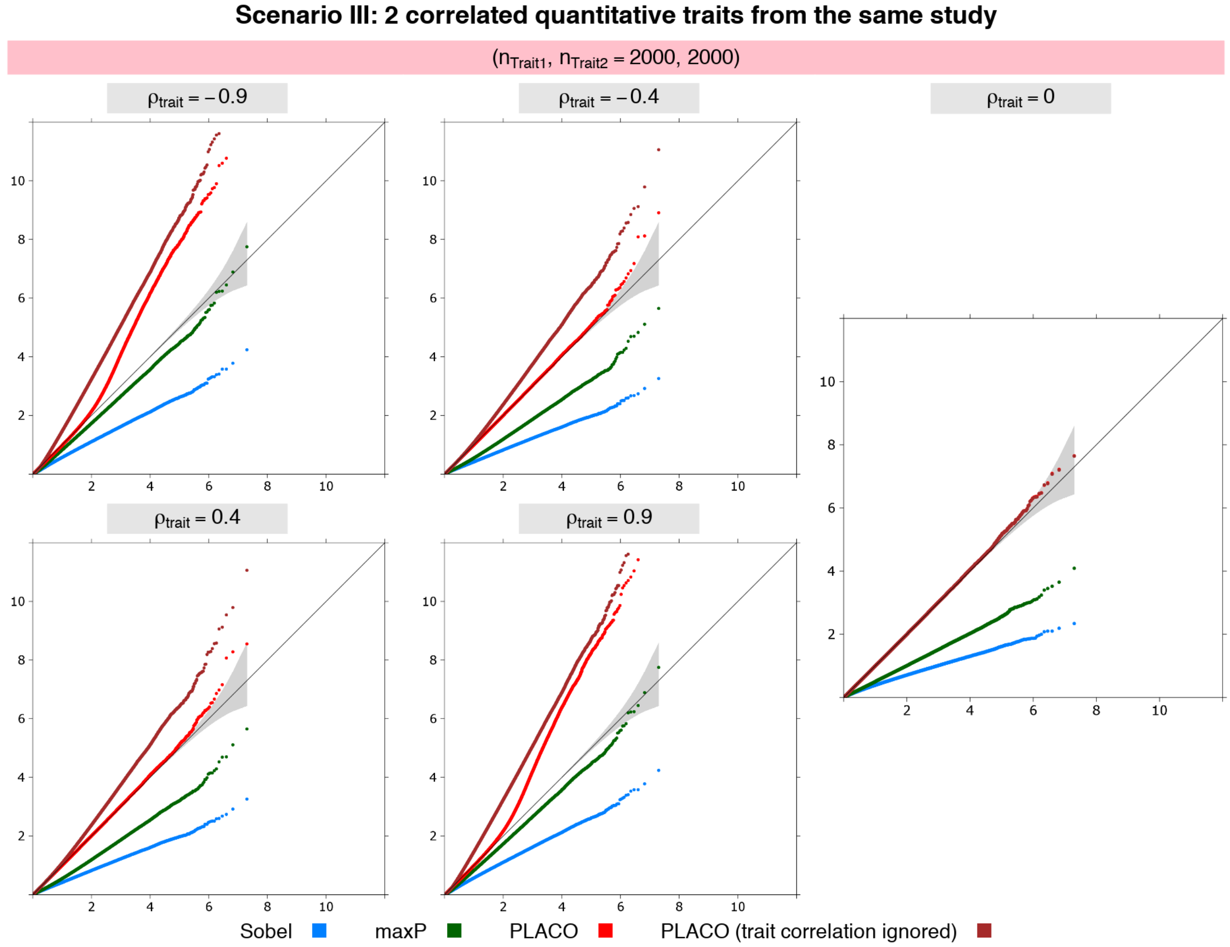


**Figure S3:** Scenario III: QQ plots for null data on 2 correlated traits where each trait is measured on the same $2,000$ individuals. Observed($-\log_{10}$p-values) are plotted on the y-axis and Expected($-\log_{10}$p-values) on the x-axis. Type I error performance of tests of pleiotropic effect of a genetic variant on the 2 traits is based on 9.99 million null variants with genetic effects that are either $\{\beta_1 = 0 = \beta_2\}$ or $\{\beta_1 = 0, \beta_2$ explains 0.1% of Trait 2 variance$\}$ or $\{\beta_1$ explains 0.1% of Trait 1 variance$, \beta_2 = 0\}$. The gray shaded region represents a conservative 95% confidence interval for the expected distribution of p-values. P-values $\geq 10^{-12}$ are shown here.

#### Supplementary S4

**Type I error performance under more general simulations.** To evaluate sensitivity (if any) of type I error control of PLACO when the genetic effects under no pleiotropy (a composite null hypothesis) are not fixed at a particular non-zero value, we additionally consider a more general situation where a distribution is assumed for one of the genetic effects. We use a normal distribution with mean 0 and standard deviation 0.1 for the genetic effect of the first trait (the choice of this distribution is motivated by the distribution of effect sizes of common variants across many complex human traits[3]). The genetic effect of the second trait is fixed at 0 (or odds ratio = 1). In other words, out of the 10 million genetic variants, we assume 99.9% variants to be under either the global null $H_{00}$ (i.e., none of the traits is associated) or the sub-null $H_{02}$ (i.e., only first trait is associated). For the 0.1% non-null variants, we assume the same distribution for the first genetic effect and fix the genetic effect for the second trait at an arbitrary non-null value.

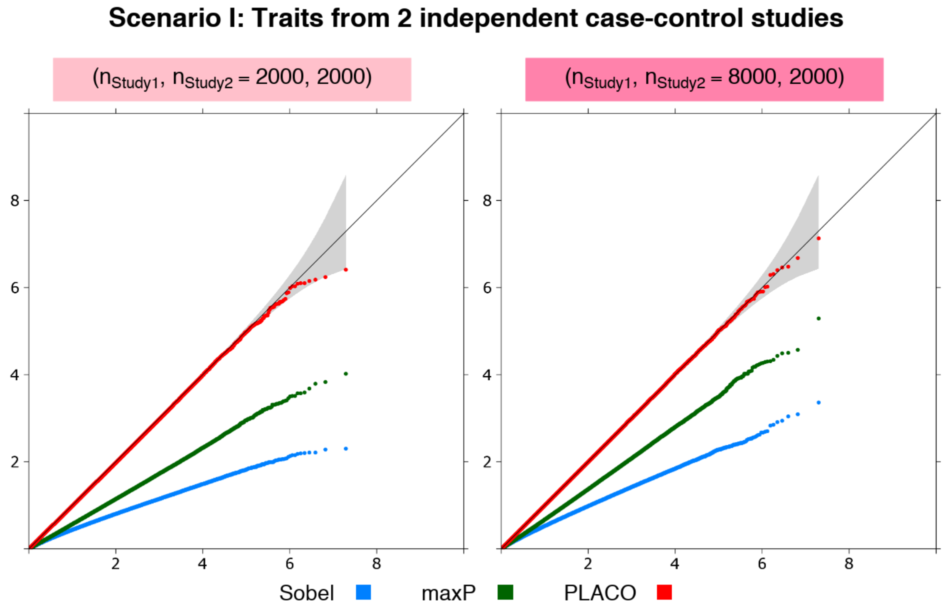


**Figure S4:** Scenario I: QQ plots for null data on traits from 2 independent case-control studies with distribution assumed for genetic effect of 1 trait. Observed($-\log_{10}$p-values) are plotted on the y-axis and Expected($-\log_{10}$p-values) on the x-axis. Either each study has 1,000 unrelated cases and 1,000 unrelated controls, or Study 1 has 4 times sample size as Study 2, where Study 2 has 1,000 unrelated cases and 1,000 unrelated controls. Type I error performance of tests of pleiotropic effect of a genetic variant on the 2 traits is based on 9.99 million null variants with genetic effects that are $\{\beta_1 \sim N(0, 0.1^2),\ \beta_2 = 0\}$. The gray shaded region represents a conservative 95% confidence interval for the expected distribution of p-values. P-values $\geq 10^{-10}$ are shown here.

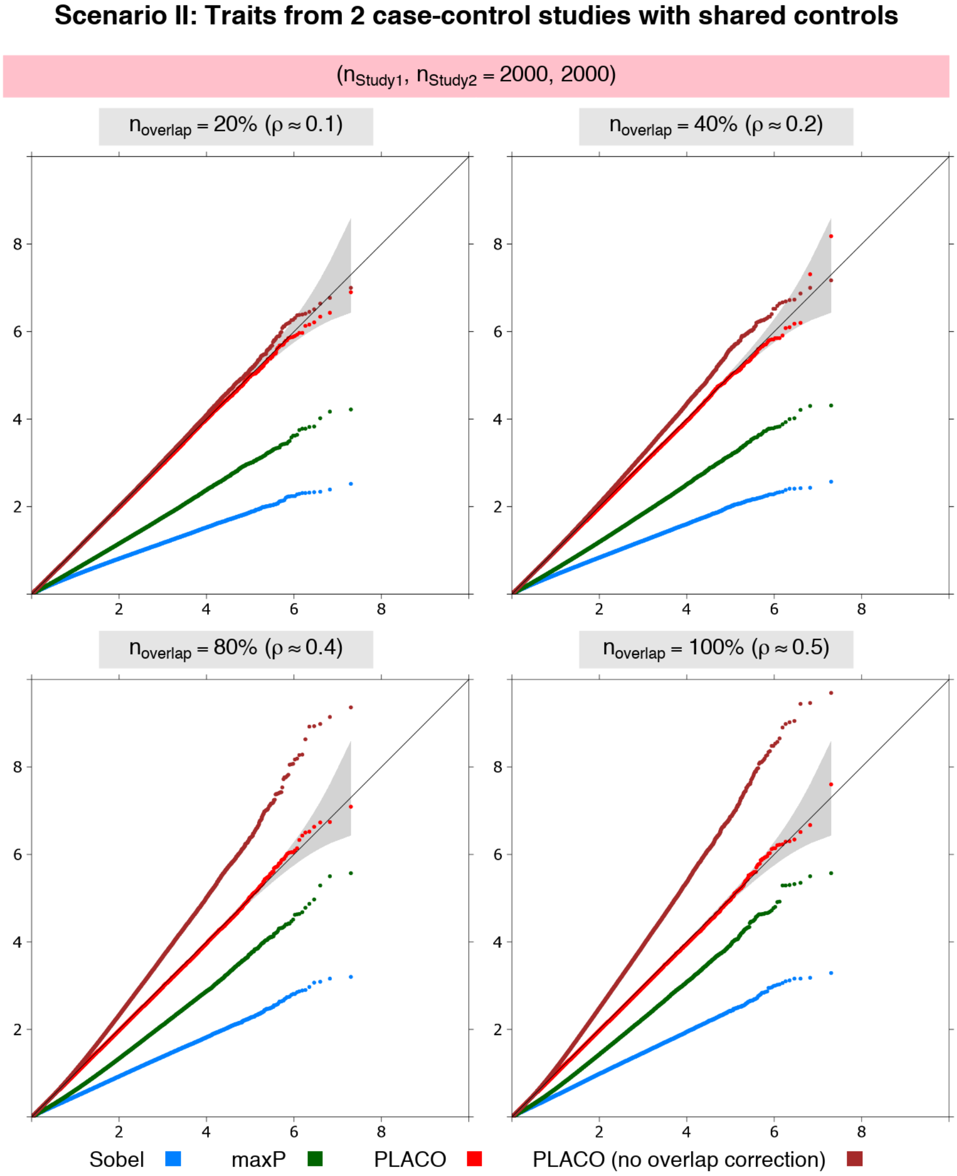


**Figure S5:** Scenario II: QQ plots for null data on traits from 2 case-control studies with different proportions of overlapping controls and with distribution assumed for genetic effect of 1 trait. Observed($-\log_{10}$p-values) are plotted on the y-axis and Expected($-\log_{10}$p-values) on the x-axis. Equal study sample size, and equal case-control size assumed in each study. Each study has $1,000$ unrelated cases and $1,000$ unrelated controls, of which either 20%, 40%, 80% or 100% of the controls are shared between the two studies. Type I error performance of tests of pleiotropic effect of a genetic variant on the 2 traits is based on 9.99 million null variants with genetic effects that are $\{\beta_1 \sim N(0, 0.1^2),\ \beta_2 = 0\}$. The gray shaded region represents a conservative 95% confidence interval for the expected distribution of p-values. P-values $\geq 10^{-10}$ are shown here.

#### Supplementary S5

##### Additional text, tables and figures for the T2D-PrCa analysis.

*More on gene-set enrichment analysis.* FUMA also performed enrichment analyses in other annotated gene sets described in Molecular Signatures Database (MSigDB v7.0)[4] and in curated biological pathways from WikiPathways[5]. We found significant enrichment in 3 gene sets representing expression signatures of genetic and chemical perturbations (Figure S14); in 2 gene sets representing potential targets of regulation by transcription factors (Figure S15); and in 1 gene set representing cell states and perturbations within the immune system (Figure S16). Delving deeper into these gene sets and pathways may provide knowledge about T2D-PrCa etiology and may shed light on the observed T2D-PrCa inverse association[6]; however delving deeper is beyond the scope of this article.

*Functional enrichment analysis.* We tested enrichment of functional consequences of 43 loci using FUMA (we excluded the MHC locus from all analyses because of strong SNP associations in this long-range and complex LD block that complicates fine-mapping efforts[7]). Fisher's exact tests of enrichment for 11 annotations show significant enrichment of SNPs in flanking regions such as 3-prime ($p_{\text{Fisher}} = 2.4 \times 10^{-23}$) and 5-prime UTRs ($p_{\text{Fisher}} = 7.2 \times 10^{-5}$) (Table S2). We found 46 (3.6%) significant SNPs spread across 19 loci have CADD scores[8] $> 12.37$ (the suggested threshold for deleteriousness, as reported in the FUMA documentation). RegulomeDB categorical scores[9] predict 35 SNPs across 9 loci to affect binding and linked to expression of a gene target, while 33 SNPs across 18 loci are likely to affect binding only. Majority of the significant SNPs has highly significant *cis*-regulatory effects ($p < 5 \times 10^{-8}$) on gene expression in whole blood from eQTLGen Consortium.

**Table S2:** Enrichment statistics for different functional consequences (annotations from ANNOVAR) of SNPs in LD with the lead SNPs from all 43 loci detected by PLACO. 1000G Phase 3 European population is used as reference panel.

| Annotation | Count (ref.) | Prop. (ref.) | Count (here) | Prop. (here) | Enrichment | $p_{\text{Fisher}}$ |
|---|---|---|---|---|---|---|
| UTR3 | 233824 | 0.00932 | 211 | 0.020 | 2.150 | $2.4 \times 10^{-23}$ |
| UTR5 | 71546 | 0.00285 | 54 | 0.005 | 1.798 | $7.2 \times 10^{-5}$ |
| downstream | 284177 | 0.01133 | 132 | 0.013 | 1.107 | $2.5 \times 10^{-1}$ |
| exonic | 254736 | 0.01016 | 131 | 0.012 | 1.225 | $2.2 \times 10^{-2}$ |
| intergenic | 11684523 | 0.46586 | 2252 | 0.214 | 0.459 | 0 |
| intronic | 9137749 | 0.36432 | 6308 | 0.599 | 1.645 | 0 |
| ncRNA_exonic | 259951 | 0.01036 | 106 | 0.010 | 0.972 | $8.1 \times 10^{-1}$ |
| ncRNA_intronic | 2884355 | 0.11500 | 1211 | 0.115 | 1.000 | $9.9 \times 10^{-1}$ |
| ncRNA_splicing | 1313 | 0.00005 | 0 | 0.000 | 0.000 | 1 |
| splicing | 2830 | 0.00011 | 0 | 0.000 | 0.000 | $6.4 \times 10^{-1}$ |
| upstream | 266686 | 0.01063 | 121 | 0.011 | 1.081 | $3.9 \times 10^{-1}$ |

Annotation: Functional consequence of SNPs
Count (ref.): Number of SNPs with the corresponding annotation in the reference panel
Prop. (ref.): Proportion of SNPs with the corresponding annotation in the reference panel
Count (here): Number of SNPs with the corresponding annotation in the candidate set of SNPs from current GWAS
Prop. (here): Proportion of SNPs with the corresponding annotation in the candidate set of SNPs from current GWAS
Enrichment: Ratio of prop. (here) to prop. (ref.); value $>1$ indicates annotation is enriched, otherwise depleted
$p_{\text{Fisher}}$ : p-value from Fisher's exact test (2-sided test)

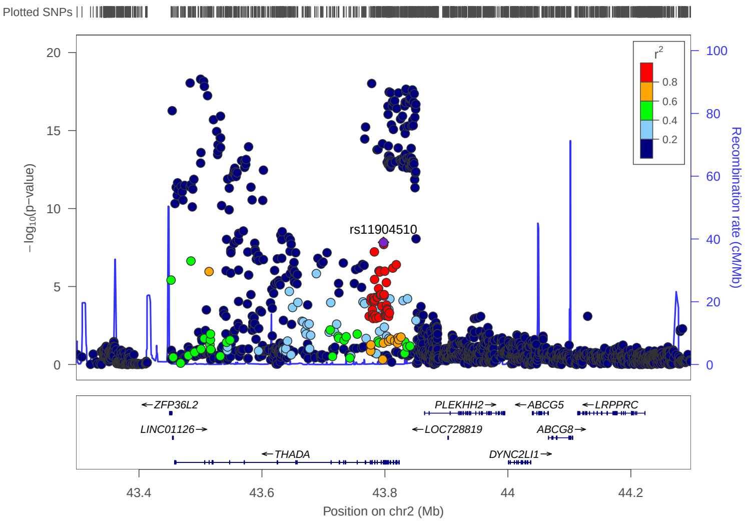


(a) T2D p-values

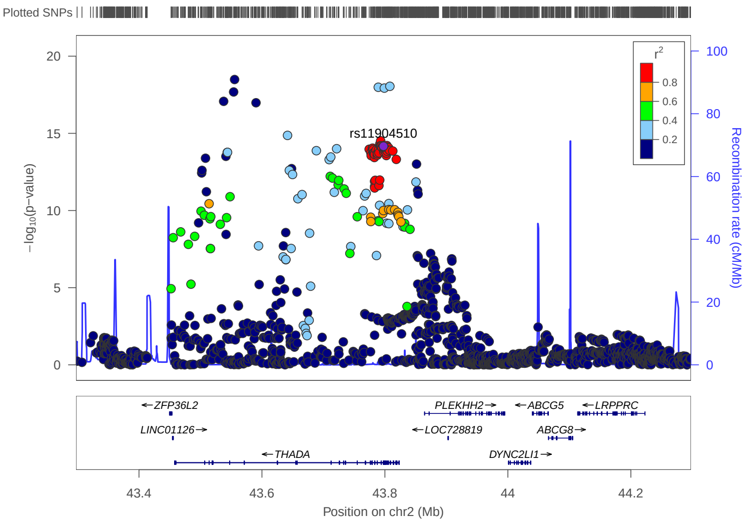


(b) PrCa p-values

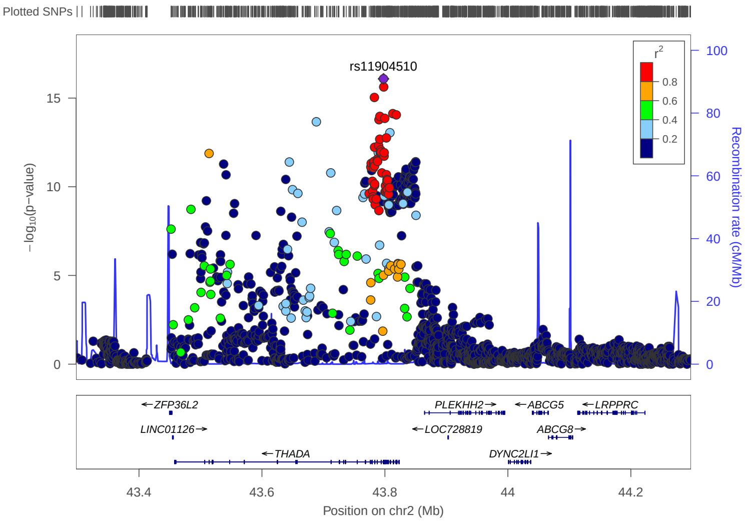


(c) PLACO p-values

**Figure S6:** Locuszoom plots of association p-values for variants in and around gene *THADA*.

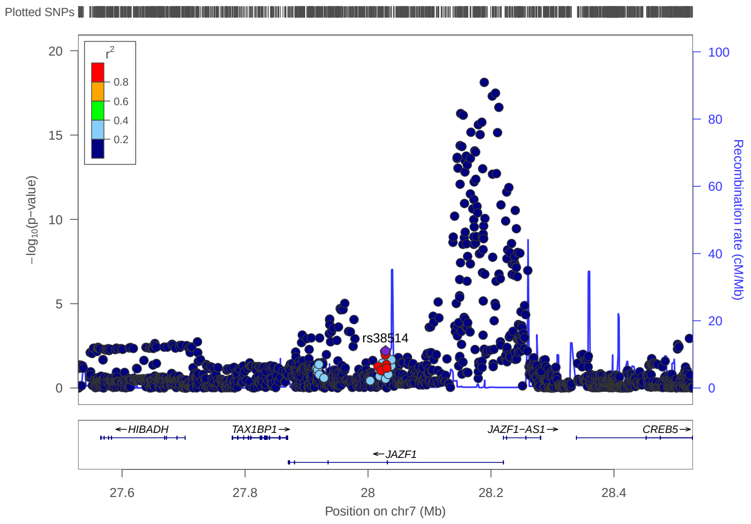


(a) T2D p-values

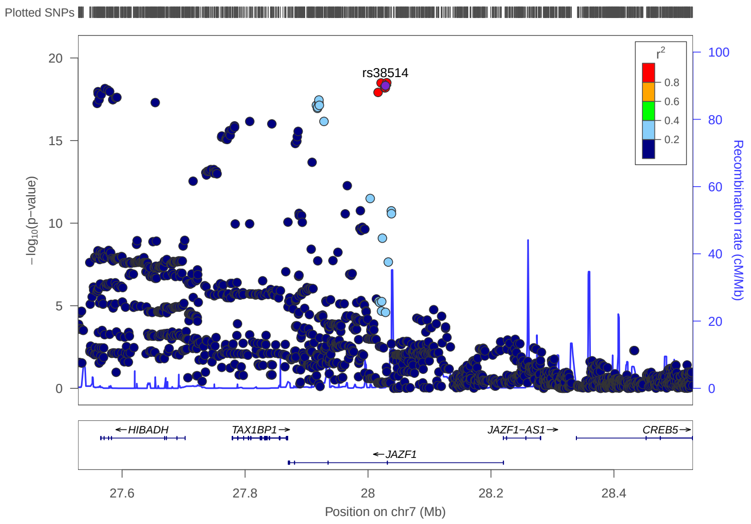


(b) PrCa p-values

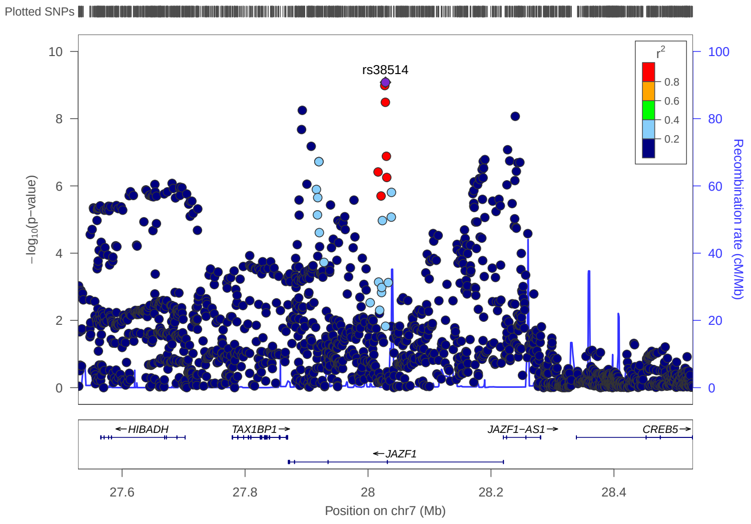


(c) PLACO p-values

**Figure S7:** Locuszoom plots of association p-values for variants in and around gene *JAZF1*.

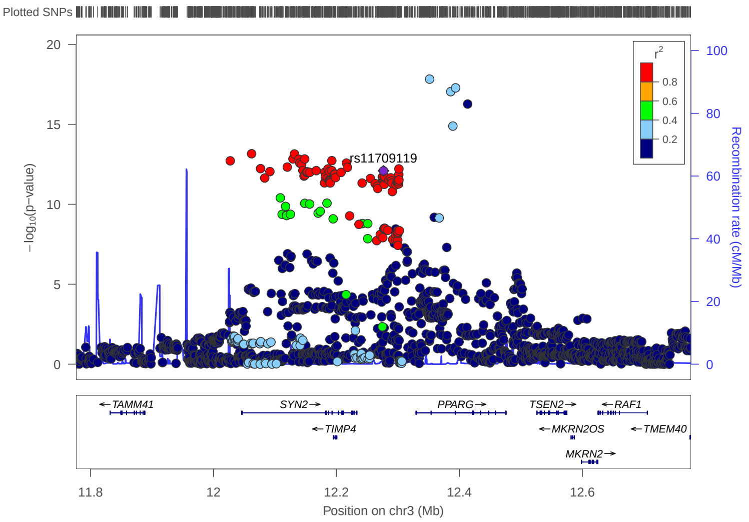


(a) T2D p-values

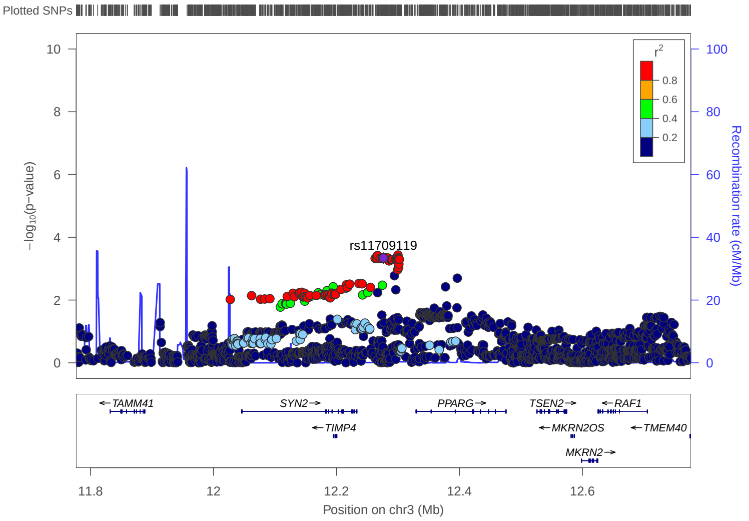


(b) PrCa p-values

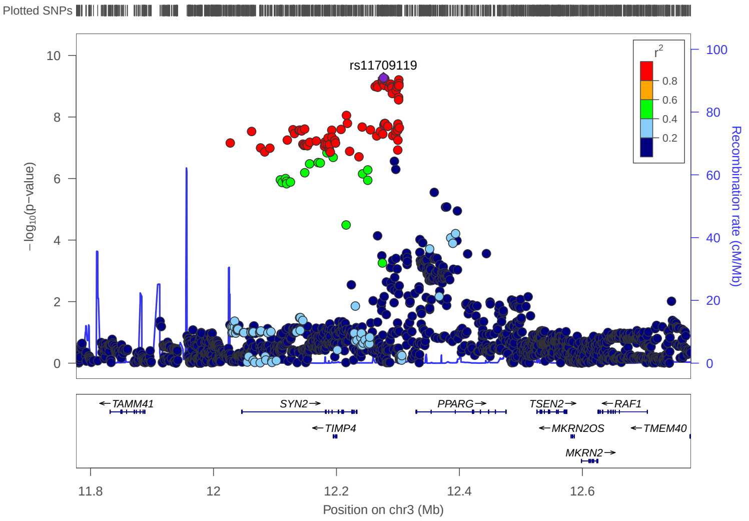


(c) PLACO p-values

**Figure S8:** Locuszoom plots of association p-values for variants in and around gene *PPARG*.

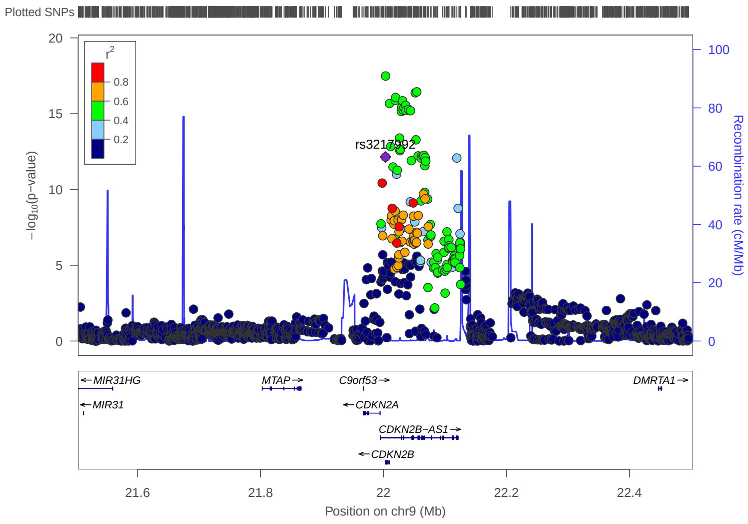


(a) T2D p-values

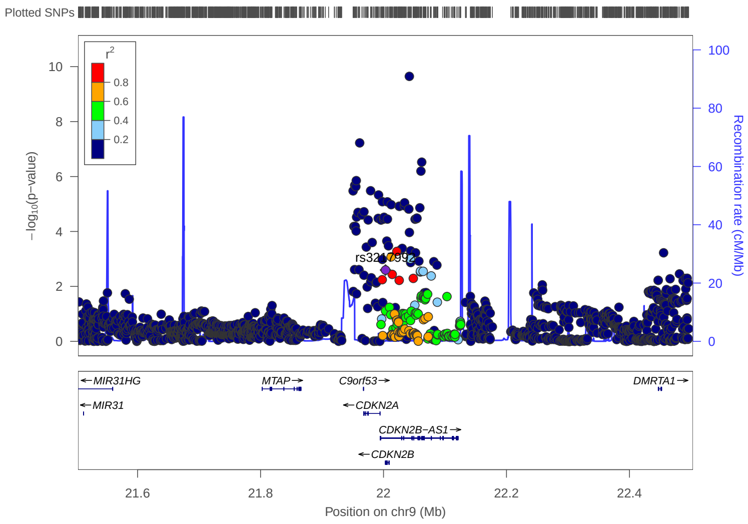


(b) PrCa p-values

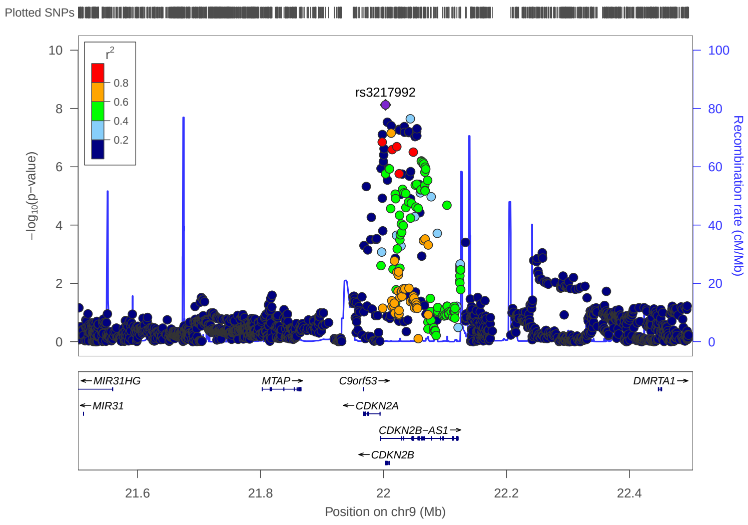


(c) PLACO p-values

**Figure S9:** Locuszoom plots of association p-values for variants in and around gene *CDKN2A*.

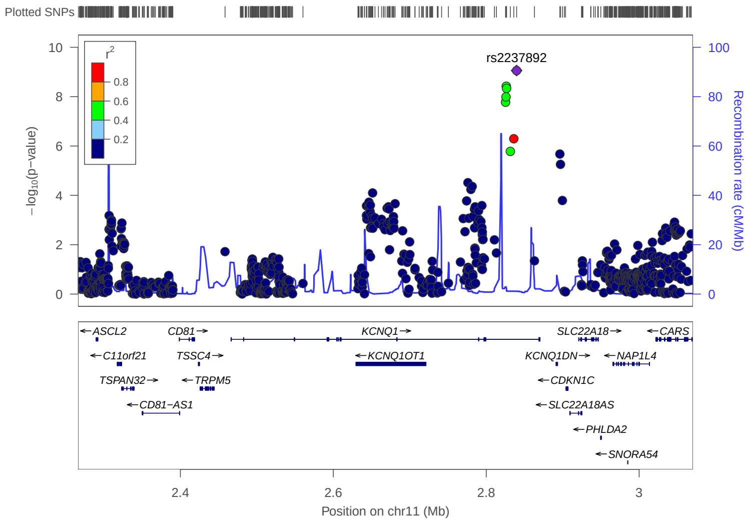


(a) T2D p-values

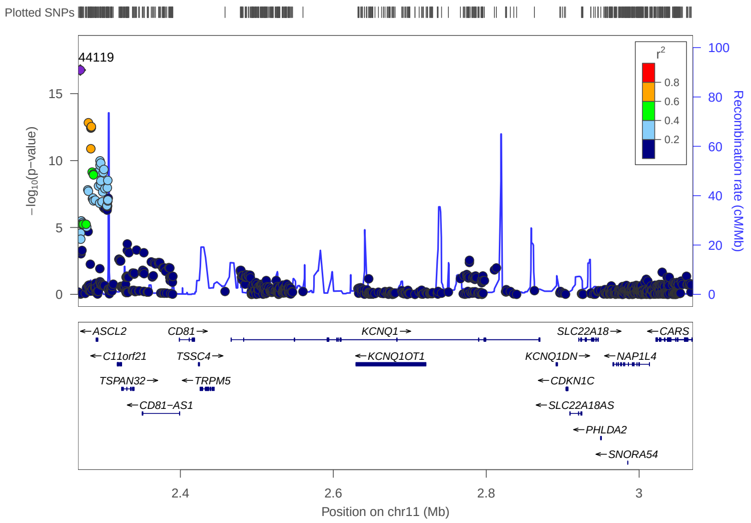


(b) PrCa p-values

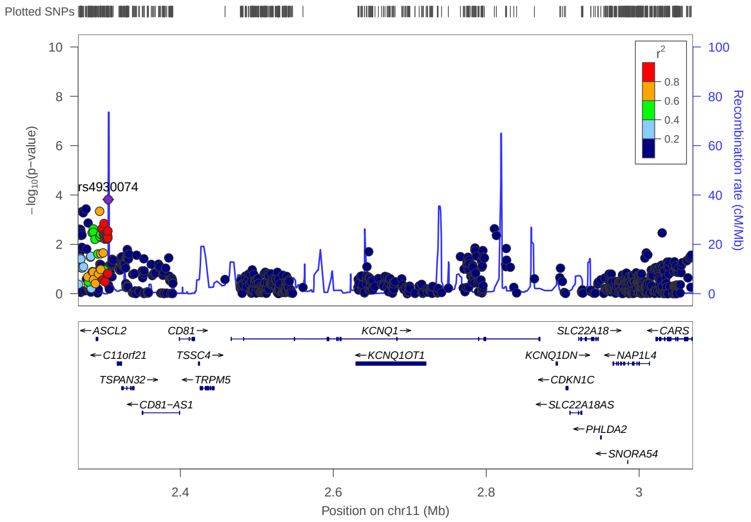


(c) PLACO p-values

**Figure S10:** Locuszoom plots of association p-values for variants in and around gene *KCNQ1*.

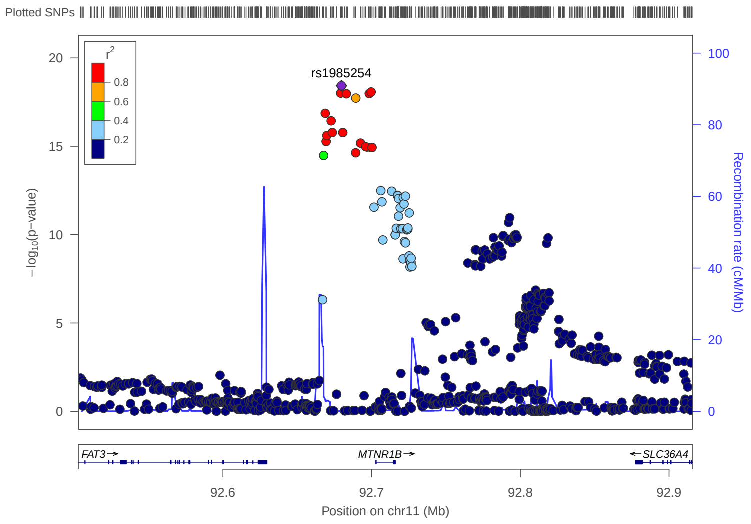


(a) T2D p-values

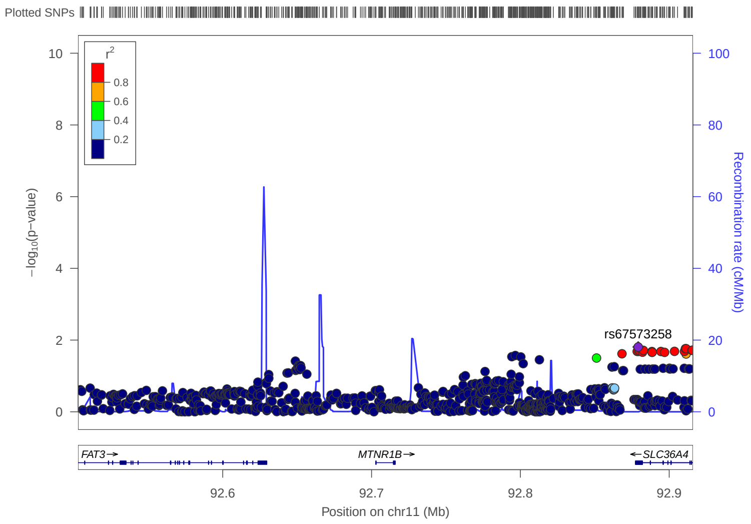


(b) PrCa p-values

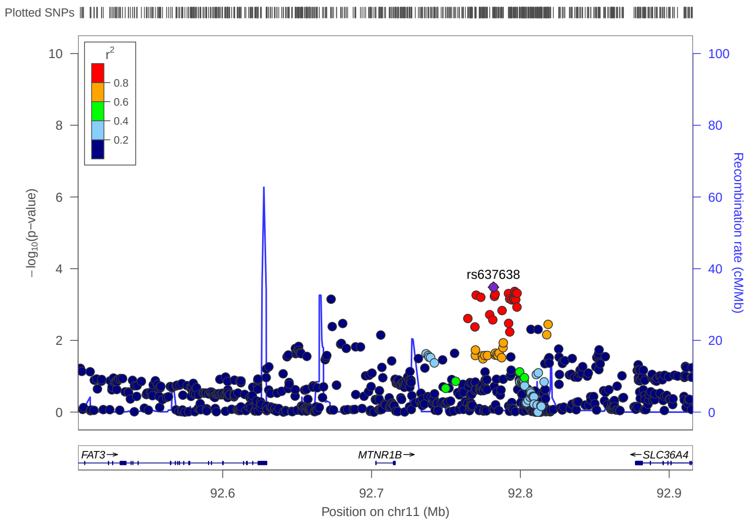


(c) PLACO p-values

**Figure S11:** Locuszoom plots of association p-values for variants in and around gene *MTNR1B*.

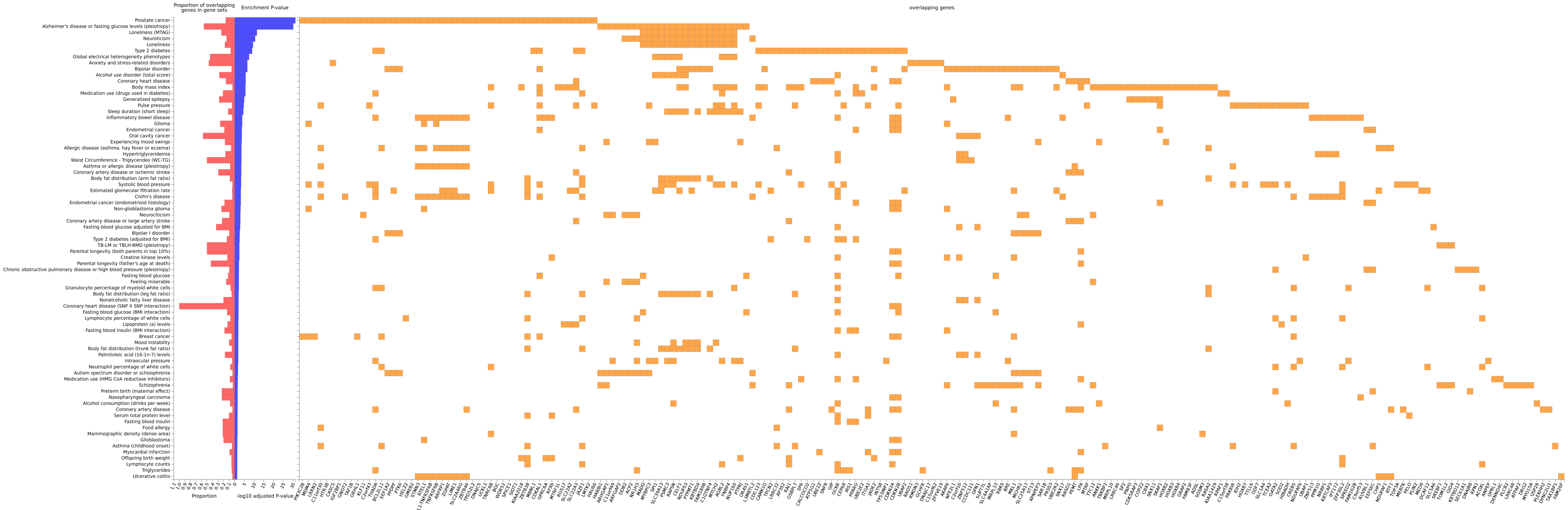


**Figure S12:** Mapped genes (as done by `FUMA`) for the 43 pleiotropic loci detected by PLACO were tested for enrichment in GWAS catalog reported genes across diseases and traits.

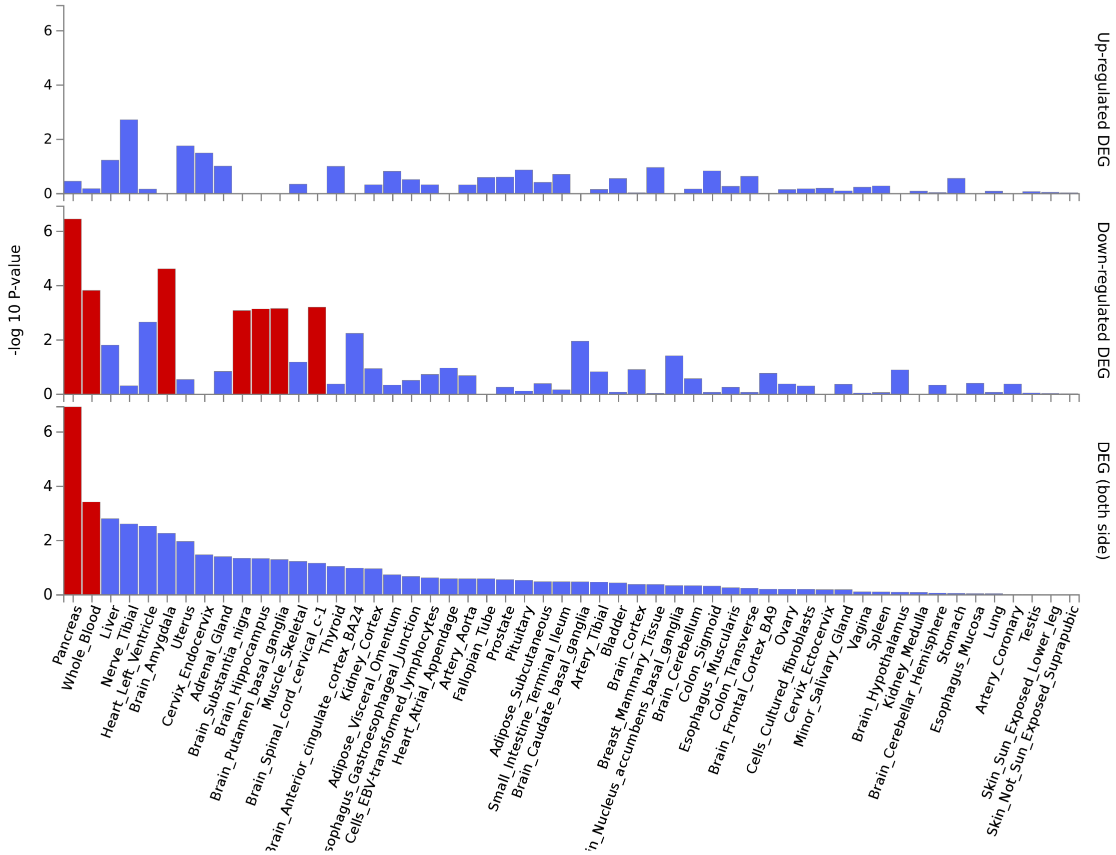


**Figure S13:** Mapped genes (as done by `FUMA`) for the 43 pleiotropic loci detected by PLACO were tested against each of the Differentially Expressed Gene (DEG) Sets pre-calculated from GTEx v8 tissue data from 53 tissue types.

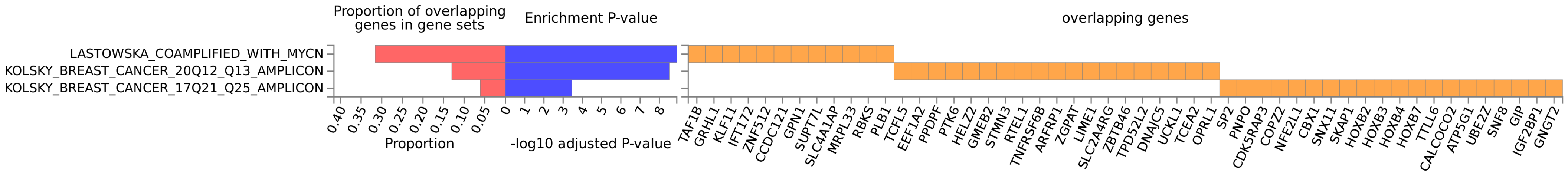


**Figure S14:** Mapped genes (as done by `FUMA`) for the 43 pleiotropic loci detected by PLACO were tested for enrichment in MsigDB C2 gene sets representing expression signatures of genetic and chemical perturbations.

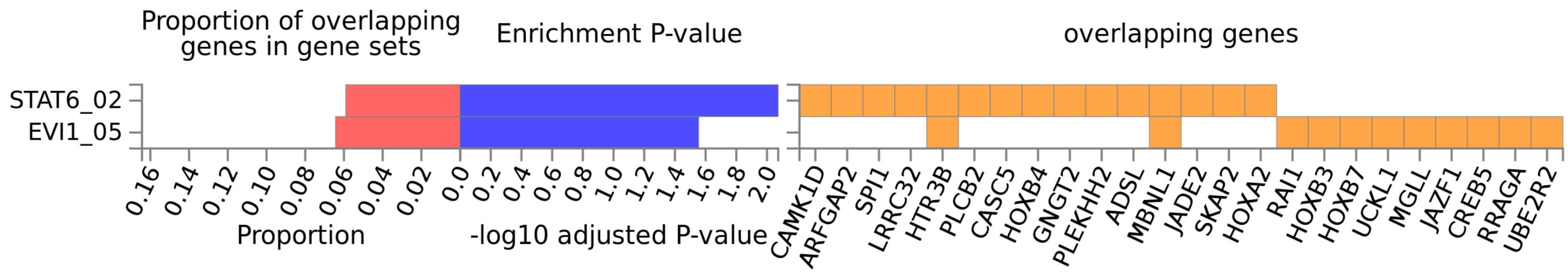


**Figure S15:** Mapped genes (as done by `FUMA`) for the 43 pleiotropic loci detected by PLACO were tested for enrichment in MsigDB C3 gene sets that share upstream cis-regulatory motifs which can function as potential transcription factor binding sites.

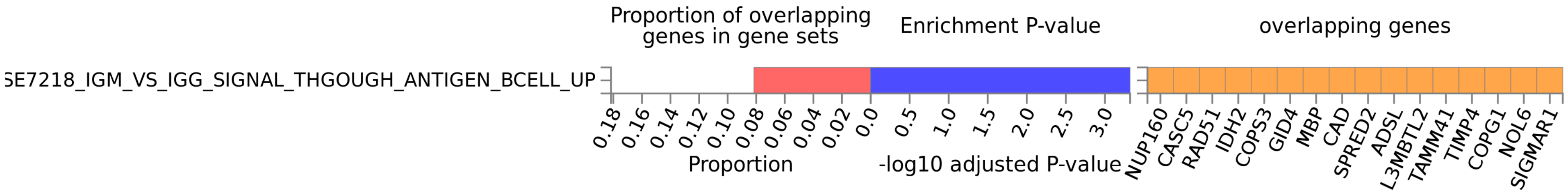


**Figure S16:** Mapped genes (as done by `FUMA`) for the 43 pleiotropic loci detected by PLACO were tested for enrichment in MsigDB C3 gene sets representing cell states and perturbations within the immune system.

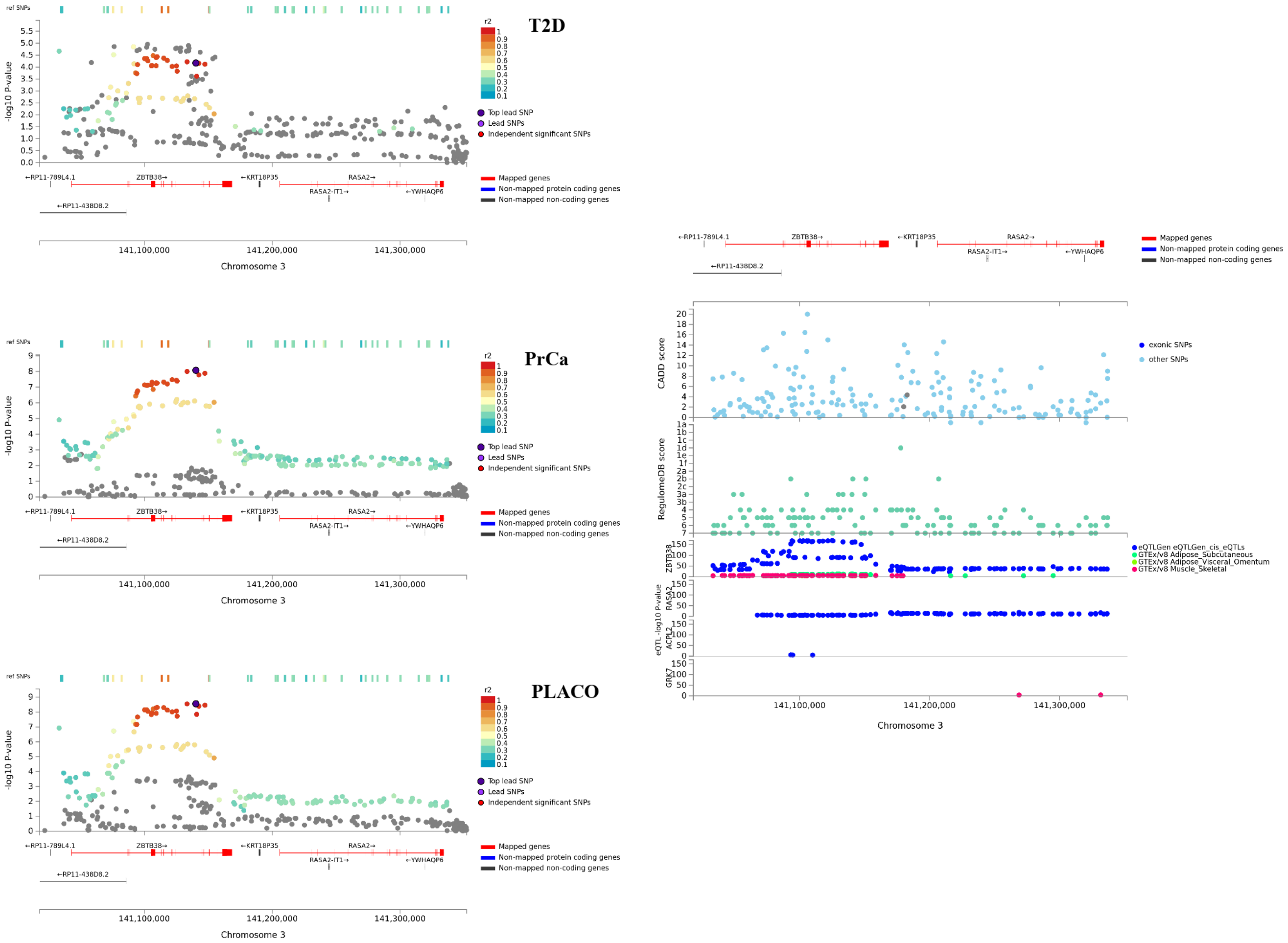


**Figure S17:** Regional association plot of significant pleiotropic locus near *ZBTB38* with annotations such as CADD scores, RegulomeDB scores, and *cis* eQTL association p-values from 6 tissues (whole blood from eQTLGen Consortium; and adipose, liver, muscle-skeletal, pancreas, and prostate tissues from GTEx v8.

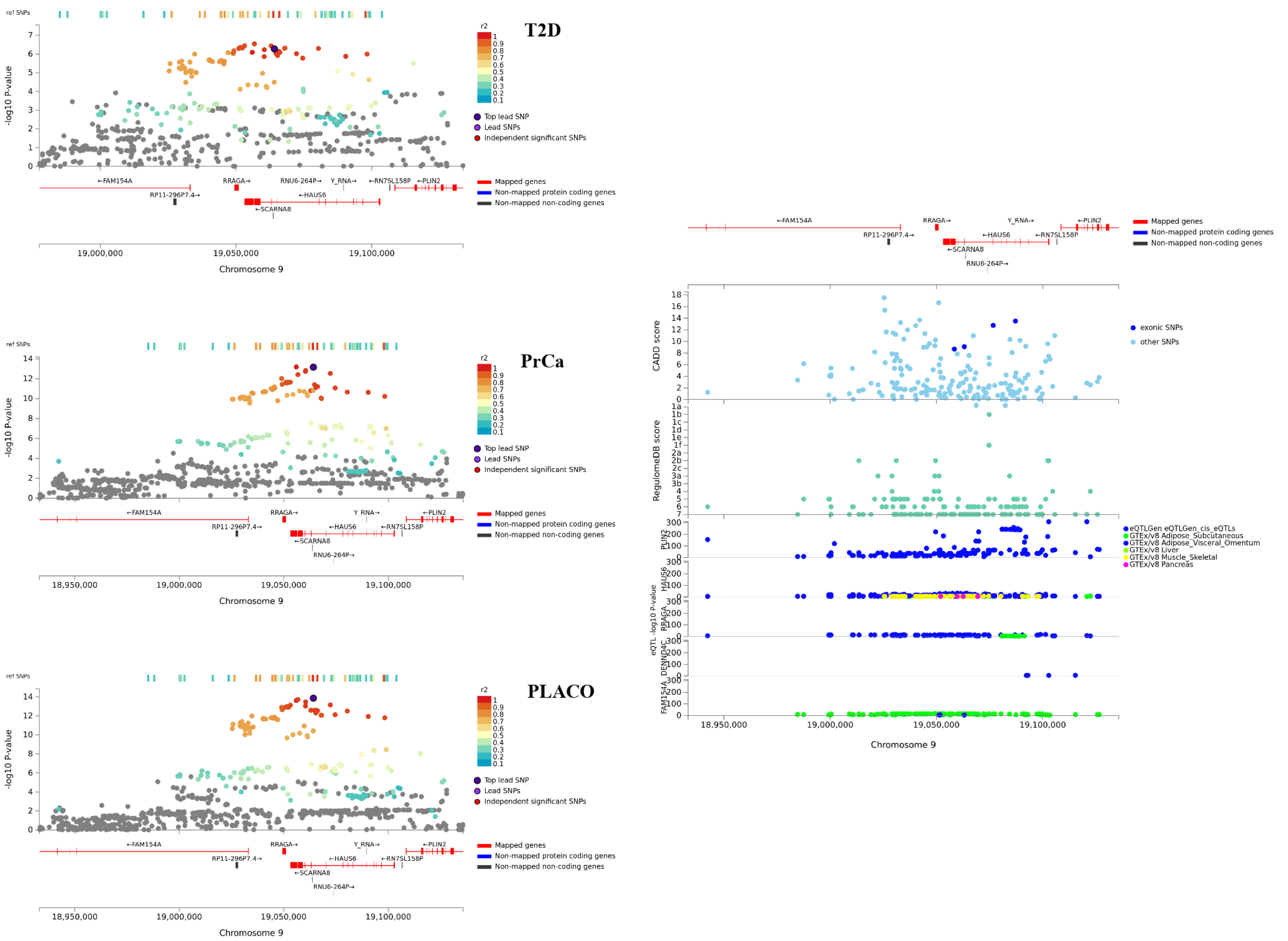


**Figure S18:** Regional association plot of significant pleiotropic locus near *HAUS6* with annotations such as CADD scores, RegulomeDB scores, and *cis* eQTL association p-values from 6 tissues (whole blood from eQTLGen Consortium; and adipose, liver, muscle-skeletal, pancreas, and prostate tissues from GTEx v8.

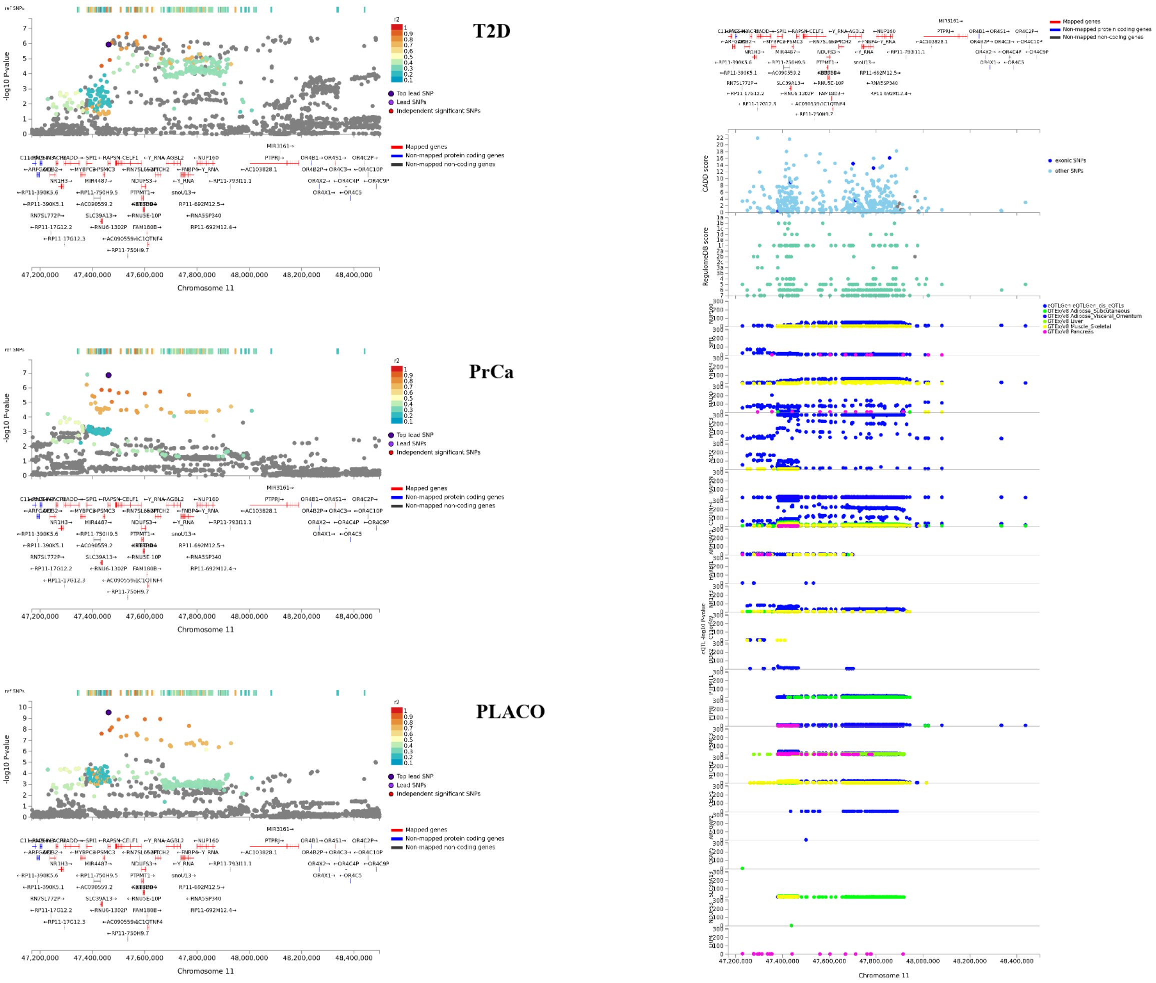


**Figure S19:** Regional association plot of significant pleiotropic locus near *RAPSN* with annotations such as CADD scores, RegulomeDB scores, and *cis* eQTL association p-values from 6 tissues (whole blood from eQTLGen Consortium; and adipose, liver, muscle-skeletal, pancreas, and prostate tissues from GTEx v8.

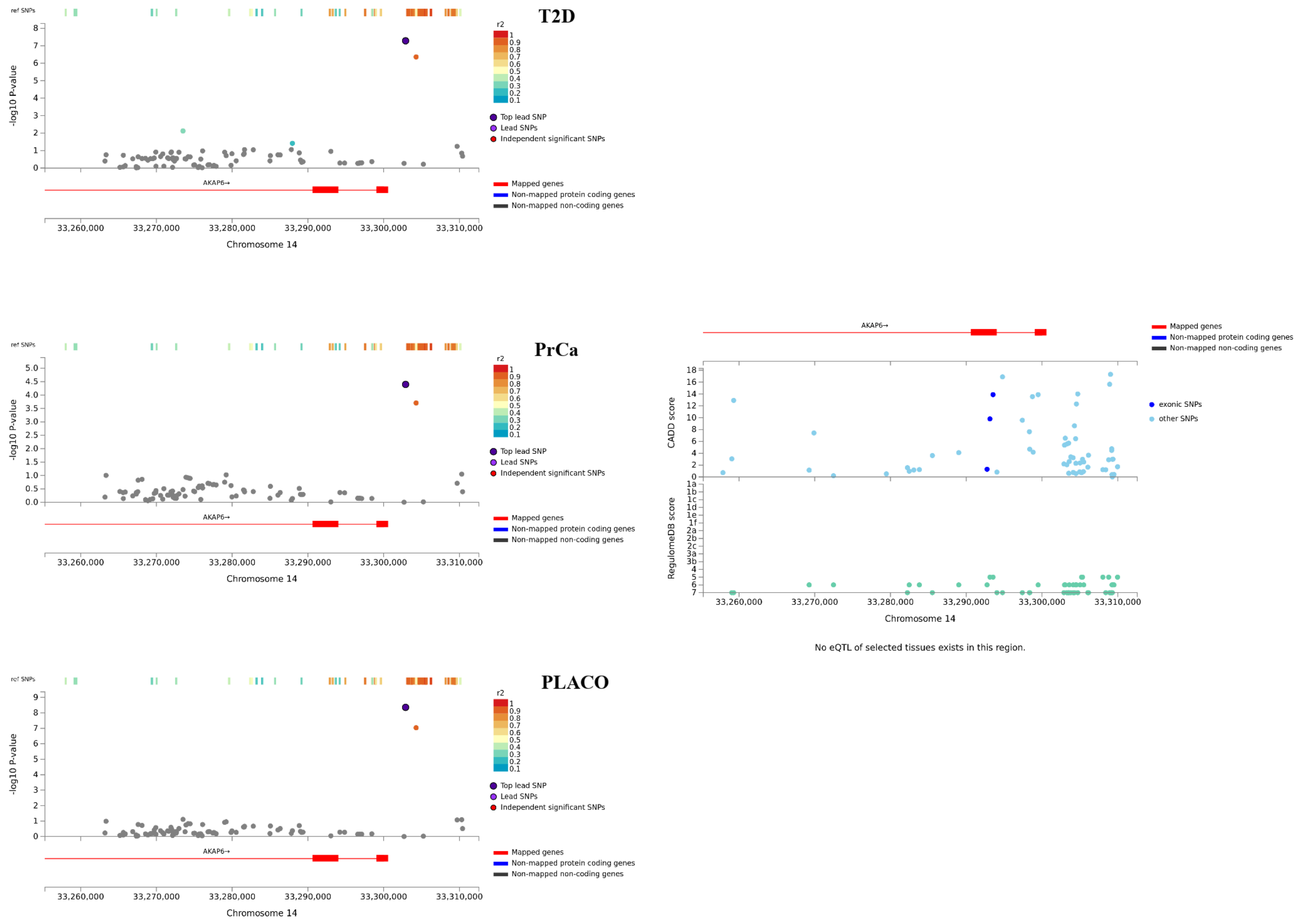


**Figure S20:** Regional association plot of significant pleiotropic locus near *AKAP6* with annotations such as CADD scores, RegulomeDB scores, and *cis* eQTL association p-values from 6 tissues (whole blood from eQTLGen Consortium; and adipose, liver, muscle-skeletal, pancreas, and prostate tissues from GTEx v8.

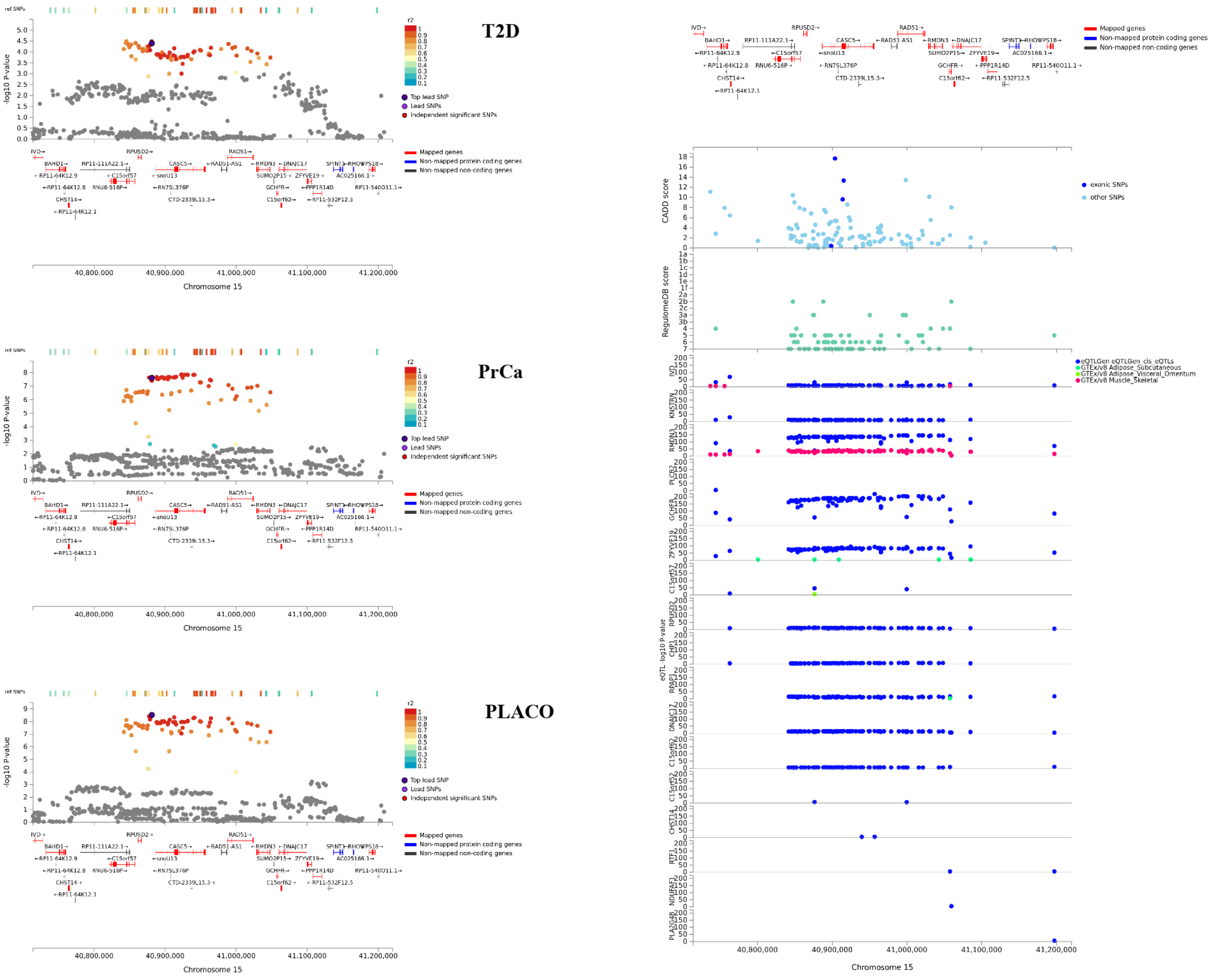


**Figure S21:** Regional association plot of significant pleiotropic locus near *KNL1* with annotations such as CADD scores, RegulomeDB scores, and *cis* eQTL association p-values from 6 tissues (whole blood from eQTLGen Consortium; and adipose, liver, muscle-skeletal, pancreas, and prostate tissues from GTEx v8.

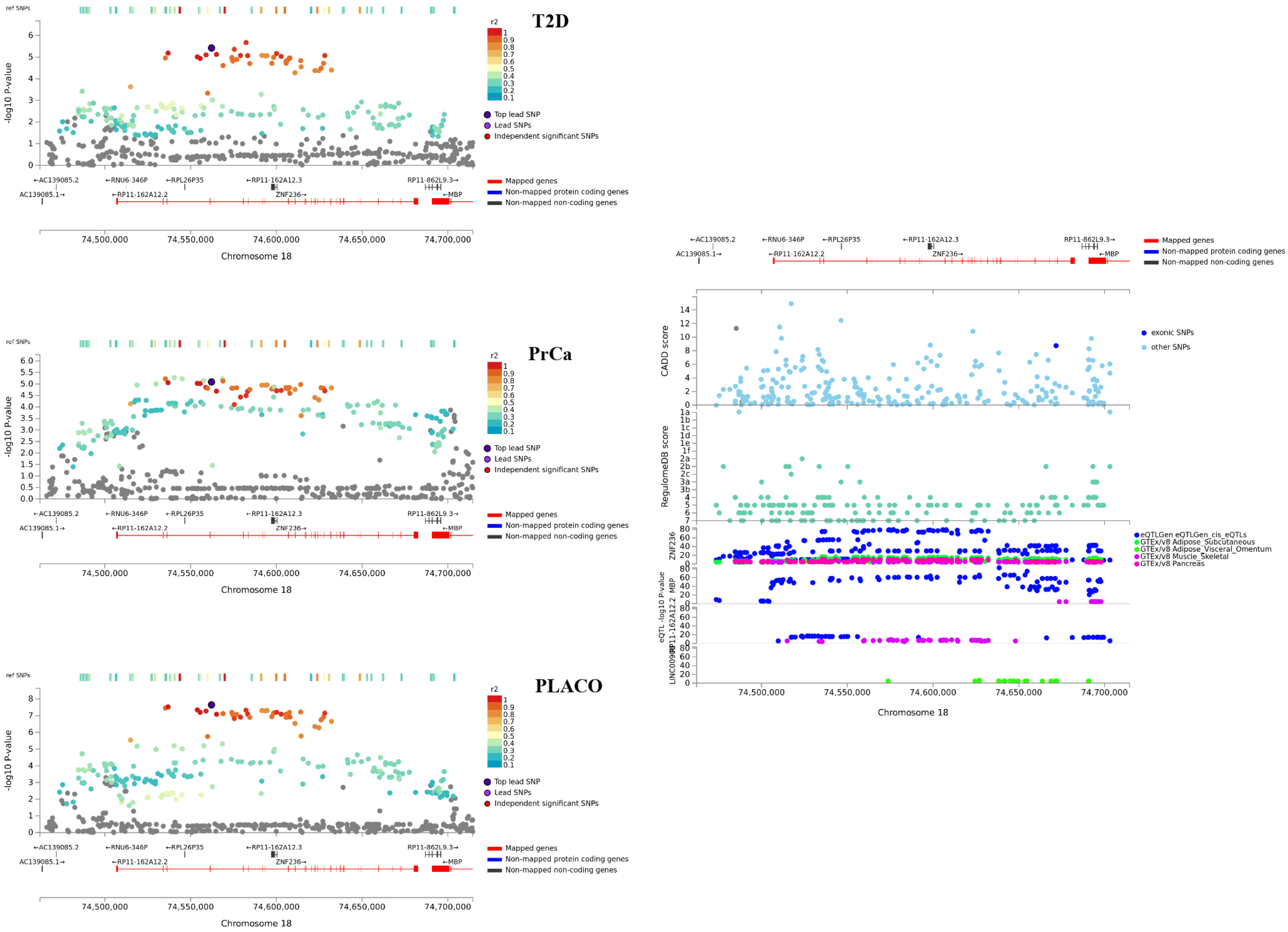


**Figure S22:** Regional association plot of significant pleiotropic locus near *ZNF236* with annotations such as CADD scores, RegulomeDB scores, and *cis* eQTL association p-values from 6 tissues (whole blood from eQTLGen Consortium; and adipose, liver, muscle-skeletal, pancreas, and prostate tissues from GTEx v8.

### References


[1] Ray, D. and Boehnke, M. Methods for meta-analysis of multiple traits using GWAS summary statistics. *Genet Epidemiol*, 42(2):134–145, 2018.

[2] Lin, D.-Y. and Sullivan, P. Meta-analysis of genome-wide association studies with overlapping subjects. *Am J Hum Genet*, 85(6):862–872, 2009.

[3] Zhang, Y., Qi, G., Park, J.-H., and Chatterjee, N. Estimation of complex effect-size distributions using summary-level statistics from genome-wide association studies across 32 complex traits. *Nat Genet*, 50(9):1318–1326, 2018.

[4] Liberzon, A., Subramanian, A., Pinchback, R., Thorvaldsdóttir, H., Tamayo, P., and Mesirov, J. P. Molecular signatures database (MSigDB) 3.0. *Bioinformatics*, 27(12): 1739–1740, 2011.

[5] Kutmon, M., Riutta, A., Nunes, N., Hanspers, K., Willighagen, E. L., Bohler, A., Mélius, J., Waagmeester, A., Sinha, S. R., Miller, R., et al. WikiPathways: capturing the full diversity of pathway knowledge. *Nucleic Acids Res*, 44(D1):D488–D494, 2015.

[6] Meyer, T. E., Boerwinkle, E., Morrison, A. C., Volcik, K. A., Sanderson, M., Coker, A. L., Pankow, J. S., and Folsom, A. R. Diabetes genes and prostate cancer in the Atherosclerosis Risk in Communities study. *Cancer Epidemiol Biomarkers Prev*, 19 (2):558–565, 2010.

[7] Mahajan, A., Taliun, D., Thurner, M., Robertson, N. R., Torres, J. M., Rayner, N. W., Payne, A. J., Steinthorsdottir, V., Scott, R. A., Grarup, N., et al. Fine-mapping type 2 diabetes loci to single-variant resolution using high-density imputation and islet-specific epigenome maps. *Nat Genet*, 50:1505–1513, 2018.

[8] Kircher, M., Witten, D. M., Jain, P., O'Roak, B. J., Cooper, G. M., and Shendure, J. A general framework for estimating the relative pathogenicity of human genetic variants. *Nat Genet*, 46(3):310–315, 2014.

[9] Boyle, A. P., Hong, E. L., Hariharan, M., Cheng, Y., Schaub, M. A., Kasowski, M., Karczewski, K. J., Park, J., Hitz, B. C., Weng, S., et al. Annotation of functional variation in personal genomes using RegulomeDB. *Genome Res*, 22(9):1790–1797, 2012.
